# Long-term population decline and recombination heterogeneity shape genomic diversity in the American alligator

**DOI:** 10.64898/2026.06.15.732480

**Authors:** Taylor Szasz-Green, John D. Konvalina, Benjamin B. Parrott, Eric A. Hoffman, Amy L. Dapper

## Abstract

The loss of genetic diversity caused by population bottlenecks can significantly impact the ability of a species to adapt to disease, natural disasters, and changing environments. Over the last 175, years, American alligator (*A. mississippiensis*) populations underwent a decline due to habitat destruction and exploitation followed by a rebound resulting from dedicated conservation efforts. Despite a current census population of millions, microsatellite data indicates a significant paucity of genetic variation within alligators. Using whole genome sequences from 19 individuals sampled from the species’ geographic range, we quantified nucleotide diversity, heterozygosity, inbreeding, demographic history, and fine-scale recombination rates. We find that American alligator genomes exhibit low nucleotide diversity and elevated homozygosity relative to many vertebrates, but these patterns are dominated by numerous short runs of homozygosity (ROHs), rather than long tracts indicative of recent inbreeding. Demographic history reconstruction based on site-frequency-spectrum analyses support a prolonged decline in effective population size beginning in the last glacial period, indicating that this reduced genetic diversity largely predates intensive human exploitation. Additionally, we uncover a highly structured recombination landscape, with recombination consistently elevated at distal chromosomal regions and suppressed across large central segments. This heterogeneity is associated with genomic spatial variation in nucleotide diversity, suggesting that the recombination landscape contributes to the persistence of homozygosity following population expansion. Together, our results highlight that demographic recovery does not necessarily equate to genetic recovery and show how long-term population history and genomic architecture continue to shape diversity in a “recovered” species.

## Introduction

Population bottlenecks, such as those experienced by American alligator (*Alligator mississippiensis*), are increasing in frequency due to anthropogenic impacts, including large-scale climatic shifts, over-hunting, and habitat destruction (Smith 1998; Holt 1990; Bellard et al. 2012; Cammen et al. 2018). These events can have lasting effects on genetic diversity even after census population sizes rebound, shaping the long-term capacity of populations to respond adaptively to environmental changes or disease (Nei et al. 1975; Hoelzel 1999; Lynch et al. 1995; Bouzat 2010; Olazcuaga et al. 2023). The rate and extent of genetic recovery following bottlenecks are shaped by key population genetic parameters - such as gene flow, selection, mutation rate, and recombination rate - which collectively influence how genetic variation is regained or lost across the genome (Charlesworth 2003; Schlichta et al. 2025). Importantly, these parameters vary widely among species and across genomic landscapes, contributing to heterogeneous patterns of recovery (Shapiro and Alm 2008; Ellegren 2014; Tigano and Friesen 2016; Stapley et al. 2017; Jiao and Yang 2021). Thus, there is a critical need to understand how demographic history, including dedicated conservation efforts, shape genome-wide patterns of genetic diversity and evolutionary potential.

The American alligator is touted as a remarkable American conservation success story (Eversole and Henke 2018). American alligators were one of the first species to be classified as endangered, following dwindling population numbers due to habitat destruction and over-harvesting (Eversole and Henke 2018). Due to effective wildlife population management strategies, including restocking from alligator farms and protective harvesting practices, American alligators were designated “fully recovered” only twenty years later and modern census population estimates range from 3°-5 million individuals (Eversole and Henke 2018). Despite the alligator’s designation as fully recovered, early studies of genetic variation in *A. mississippiensis* have consistently reported low mitochondrial and nuclear diversity, based on mitochondrial sequencing and microsatellite analyses across the species’ range (Glenn et al. 1998, 2002; Davis et al. 2002; Ryberg et al. 2002), with the first reference genome of *A. mississippiensis* confirming unusually low autosomal heterozygosity relative to other vertebrates (Green et al. 2014).

Interestingly, the low levels of genetic diversity observed in *A. mississippiensis* are not a general characteristic of crocodilian genomes. Across much of the clade, particularly among widespread tropical species, genome-wide heterozygosity is moderate to high, with the exception of a small number of taxa that have experienced extreme, recent population collapse or long-term isolation (van Asch et al. 2019; Bloor et al. 2015; Dever et al. 2002; Green et al. 2014; Lapbenjakul et al. 2017; Sharma et al. 2020; Velo-Antón et al. 2014). The American alligator is therefore unusual in exhibiting persistently low heterozygosity despite a broad geographic distribution and a contemporary census population numbering in the millions (Glenn et al. 2002; Davis et al. 2002; Green et al. 2014; Eversole and Henke 2018). This pattern likely reflects the species’ distinctive demographic history. In addition to a well-documented recent bottleneck driven by overharvesting and habitat loss in the nineteenth and twentieth centuries (Eversole and Henke 2018; Elsey et al. 2019), *A. mississippiensis* occupies the northernmost range of any extant crocodilian and has experienced repeated population contractions during Pleistocene glacial cycles, when cooler climates restricted populations to southern coastal refugia (Yokoyama et al. 2000; Waltari et al. 2007; Green et al. 2014). Together, this raises the question of whether the current paucity of genetic diversity observed in *A. mississippiensis* reflects recent inbreeding associated with modern management practices, or instead results from deeper demographic history predating human exploitation. Disentangling these alternatives requires genome-wide data capable of resolving both the magnitude and temporal origin of homozygosity.

Runs of Homozygosity (ROHs) - long, contiguous stretches of DNA that are identical by descent (IBD) (Ceballos et al. 2018; Shafer and Kardos 2025) - provide a powerful framework for distinguishing recent consanguineous inbreeding from more ancient population declines, as ROH length is inversely related to the time since a common ancestor (Ceballos et al. 2018; Kyriazis et al. 2025). Long ROHs reflect recent inbreeding (i.e., within the last 25 generations), while numerous short ROHs point to repeated historical population contractions (Kyriazis et al. 2025). In reptiles, extreme cases of recent bottlenecks—such as the critically endangered Chinese alligator (*Alligator sinensis*)—are characterized by extensive genome coverage by long ROHs (Yang et al. 2023; Pan et al. 2025), providing a comparative framework for interpreting patterns in *A. mississippiensis*.

Meiotic recombination further shapes genomic recovery following population bottlenecks by breaking up linkage blocks and creating new combinations of alleles (Hill and Robertson 1966; Charlesworth et al. 1993; Peñalba and Wolf 2020; Ahrens et al. 2026). Recombination rates vary widely among vertebrates and are influenced by chromosomal architecture (Stapley et al. 2017; Dumont et al. 2009; Johnston et al. 2016). Although fine-scale recombination landscapes are poorly characterized in reptiles, cytogenetic and linkage-map data from snake and lizard genomes suggest that recombination rates may be generally low and biased towards the distal regions of chromosomes in this clade, which would further slow recovery following genetic bottlenecks (Lisachov et al. 2017, 2019; Schield et al. 2020). However, fine-scale recombination landscapes in chickens, a closer relative to crocodilians, do not reveal distally biased patterns (Weng et al. 2019). Thus, understanding how recombination is distributed across alligator chromosomes is essential for interpreting patterns of diversity and homozygosity across the genome.

Here, we use whole-genome sequencing from 19 American alligators sampled across the species’ range to improve the resolution of genome-wide nucleotide diversity, characterize the frequency and distribution of runs of homozygosity (ROHs), estimate historical changes in effective population size (*N_e_*) and construct a fine-scale recombination landscape. By integrating inferences from multiple lines of fine-scale genomic information, we assess whether the reduced genetic diversity observed in *A. mississippiensis* primarily reflects recent inbreeding or deeper historical population decline. In addition, we present the first fine-scale, genome-wide recombination map for a crocodilian, extending beyond previous pedigree-based linkage maps (Miles et al. (2009)) and providing new insight into how chromosome-scale recombination structure may shape genetic diversity in this recovered species.

## Results

### Sampling & Relatedness

We sequenced whole genomes from 19 American alligators sampled from across the species’ geographic range (Supplementary Figure 1). Because including close relatedness (1*^st^ −* 2*^nd^* degree) can interfere with estimates of genetic diversity and inbreeding, we first evaluated kinship among our sampled individuals. We identified the kinship coefficient between each genome using KING software (–cluster, –degree 4) (Manichaikul et al. 2010). We inferred two groups of three individuals with third degree relationships (first cousin, great-grandparent, great-aunt/uncle, great-niece/nephew, great-grandchild, half-aunt/uncle). We also inferred two groups of individuals with fourth degree relationships (Supplementary Table 3). A map of these relationships approximate the geographic range of the species (Supplementary Table 1, Supplementary Figure 2). Importantly, the mean kinship coefficient was negative (Supplementary Table 3, *φ_Mean_* = *−*0.3065*, φ_Median_* = *−*0.3292*, φ_SD_* = 0.2230), suggesting that inferences of actual relatedness may be due to low overall patterns of genetic diversity (Fujisaki et al. 2014). This result was consistent with our expectations, as alligators generally live within a 30km radius of territory (Fujisaki et al. 2014) and most individuals represented in this study were harvested in different waterways (Supplementary Table 1).

### Nucleotide Diversity & Heterozygosity

Nucleotide diversity measures polymorphism as the average difference in nucleotides per site between two randomly selected individuals within a population (Nei and Li 1979). Genome-wide nucleotide diversity was uniformly low across all 16 autosomes, with a mean *π* of 8.80 *×* 10*^−^*^4^ (Figure 1). Nucleotide diversity showed modest spatial variation along chromosomes but was consistently elevated toward distal chromosomal regions (Figure 1). Although *π* was several-fold higher than that reported for the critically endangered Chinese alligator (*π* = 1.47 *×* 10^−4^) (Yang et al. 2023; Pan et al. 2025), it is an order of magnitude lower than values observed in most avian populations (indigenous junglefowl: *π* = 3.34 *×* 10*^−^*^3^, collared flycatcher: *π* = 3.84 *×* 10*^−^*^3^ *−* 4.01 *×* 10*^−^*^3^, and broiler chickens: *π* = 3.1 *×* 10*^−^*^3^) (Dutoit et al. 2017; Nie et al. 2019)).

**Figure 1:**
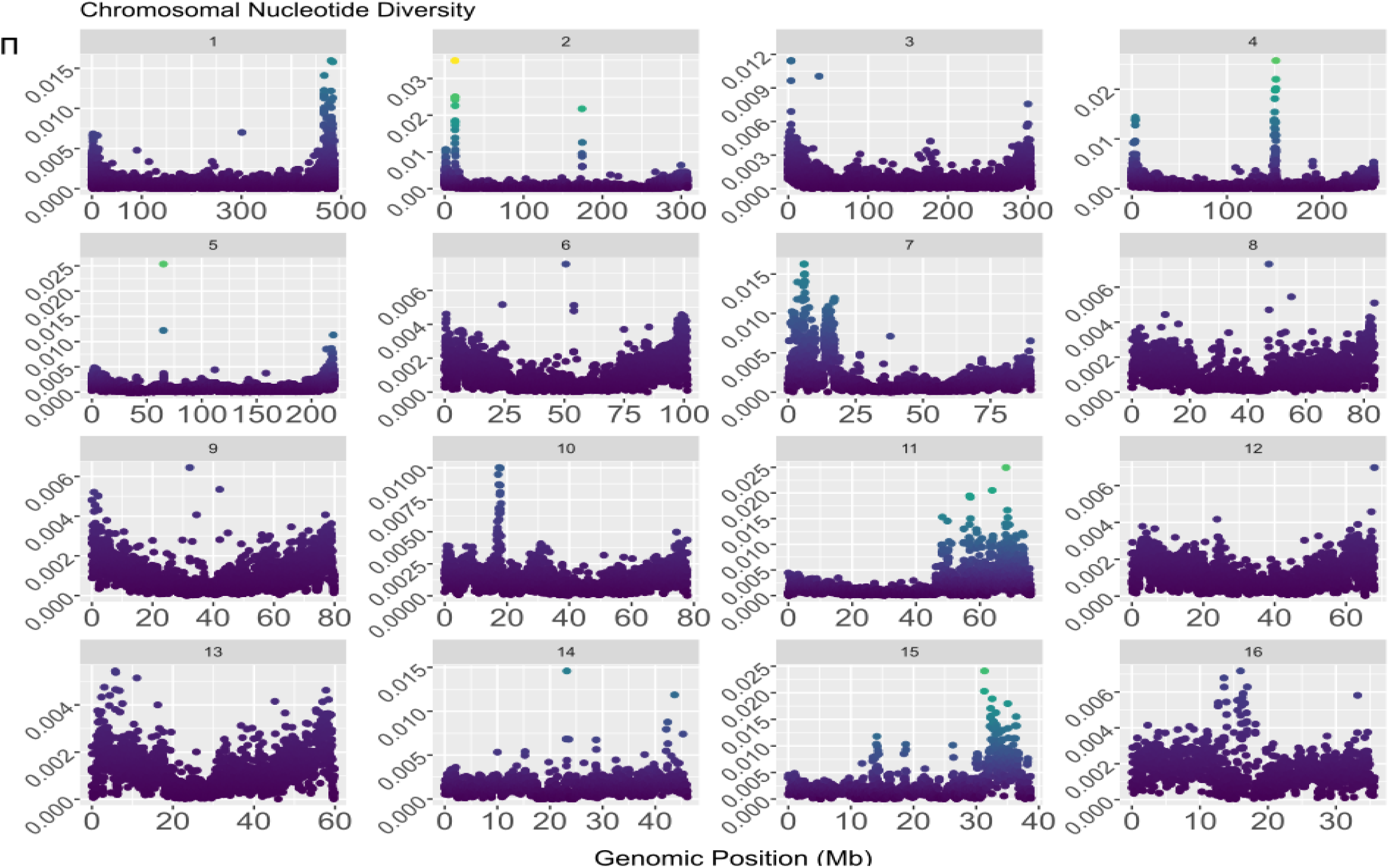
Nucleotide diversity (*π*) distribution for 16 alligator autosomes.

Because nucleotide diversity reflects differences accumulated across the genome, we next examined heterozygosity, which measures the proportion of sites at which alleles differ within individuals and more directly reflects realized genetic variation. Genome-wide autosomal heterozygosity was similarly low (*H* = 2.5*×*10*^−^*^3^), but an order of magnitude higher than than earlier microsatellite-based estimates for American alligators (*H* = 1.0 *×* 10*^−^*^4^) (Green et al. 2014) and both wild and captive populations of Chinese alligators (*H* = 1.20 *×* 10*^−^*^4^*, H* = 1.90 *×* 10*^−^*^4^ respectively) (Pan et al. 2025). When considering only polymorphic sites, there is a significant difference between observed heterozygosity (*H_O_* = 0.231) and expected heterozygosity (*H_E_* = 0.287) between individuals (*p <* 0.0001, two-tailed t-test) (Table 2), indicating an excess of homozygosity at variable loci. Our genome-wide estimates fall within the range of *H_O_* previously reported from microsatellite data (0.231 - 0.865), but is significantly lower than the mean (*H_O_* = 0.466) (Glenn et al. 1998). The expected heterozygosity is also markedly lower than seen in microsatellite data from the Chinese alligator (*H_E_*= 0.47), although differences in marker type and allele number may contribute to this discrepancy (Pan et al. 2019). Together, these results suggest that American alligator genomes are characterized by reduced standing genetic variation which is not confined to a small subset of chromosomes or individuals.

### Inbreeding & Runs of Homozygosity

To quantify genome-wide excess homozygosity, we first estimated method-of-moments inbreeding coefficients (*F_IT_*), which summarize departures from Hardy–Weinberg expectations and captures homozygosity arising from both historical and recent demographic processes. Method of moments estimates of inbreeding were consistently positive across individuals, indicating genome-wide excess homozygosity (*F_IT_*)(Table 2, *Mean* = 0.195*, Median* = 0.194*, SD* = 0.102*, Min* = 0.048*, Max* = 0.434). Because inbreeding estimates can be sensitive to estimator choice, we also calculated a genome-wide heterozygosity-based inbreeding coefficient (*F*^^^), which yielded comparable values (*Mean* = 0.141*, Median* = 0.160*, SD* = 0.086). Genome-wide *F_IT_* values are similar in magnitude to those previously reported from microsatellite data comparing Louisiana and Florida alligator populations (*F_IT_*= 0.164) (Glenn et al. 1998). Principal component and clustering analysis did not reveal evidence of strong population stratification among individuals, although the detection of subtle structure may be limited by sample size.

To resolve the genomic distribution and temporal context of homozygosity, we next characterized runs of homozygosity (ROHs), which represent contiguous tracts of identical-by-descent sequence and retain information about the timing of shared ancestry through their length distributions. Across individuals, we identified 23,279 ROHs, with a mean of 1225 segments per individual (*Median* = 1186*, SD* = 222.893) (Table 1). ROHs were classified into four length classes corresponding to progressively more recent shared ancestry (generations to most recent common ancestor (MRCA)): 0.25 - 0.5 Mb (< 200), 0.5 - 1Mb (< 100), 1Mb - 2Mb (< 50), and 2 - 4 Mb (< 25) (Figure 2C). Total genome coverage by ROHs across individuals ranged from 314.8 Mb to 750.4 Mb (*Mean* = 500.95*, Median* = 471.96*, SD* = 112.14)(Figure 2B,C). Mean ROH length for segments 250kb (*ROH*_250_) was 408.91 Kb (*Median* = 399.22*, SD* = 30.44).

**Figure 2:**
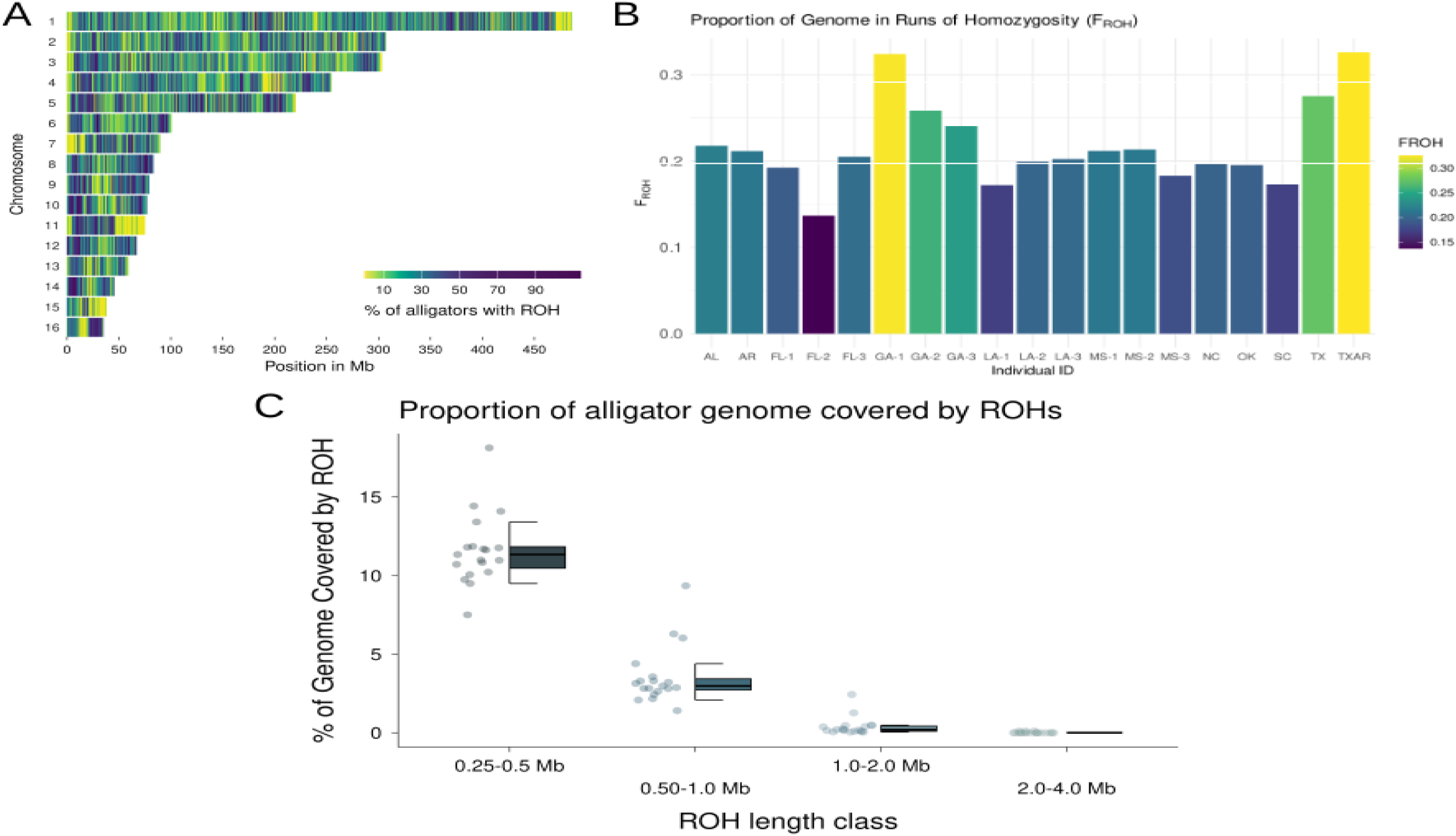
(A) Coverage of genome by runs of homozygosity (ROHs). (B) Distribution of *F_ROH_* between individuals. (C) Distribution of ROH lengths between individuals.

**Table 1:** Number of ROH segments, total genome coverage, average ROH size, and FROH per individual.

| ID | NSEG | KB | KBAVG | MB_Coverage | MB_Average | FROH |
| --- | --- | --- | --- | --- | --- | --- |
| AL | 1230 | 5.011e+5 | 407.362 | 501.055 | 0.407 | 0.218 |
| AR | 1261 | 4.869e+5 | 386.148 | 486.933 | 0.386 | 0.212 |
| FL-1 | 1107 | 4.425e+5 | 399.720 | 442.49 | 0.400 | 0.192 |
| FL-2 | 836 | 3.148e+5 | 376.564 | 314.808 | 0.377 | 0.137 |
| FL-3 | 1194 | 4.720e+5 | 395.278 | 471.962 | 0.395 | 0.205 |
| GA-1 | 1493 | 7.458e+5 | 499.552 | 745.832 | 0.500 | 0.324 |
| GA-2 | 1255 | 5.944e+5 | 473.660 | 594.443 | 0.474 | 0.259 |
| GA-3 | 1338 | 5.538e+5 | 413.916 | 553.82 | 0.414 | 0.241 |
| LA-1 | 1019 | 3.961e+5 | 388.702 | 396.087 | 0.389 | 0.172 |
| NC | 1107 | 4.554e+5 | 411.411 | 455.432 | 0.411 | 0.198 |
| MS-1 | 1238 | 4.875e+5 | 393.801 | 487.526 | 0.394 | 0.212 |
| LA-2 | 1162 | 4.587e+5 | 394.722 | 458.667 | 0.395 | 0.199 |
| OK | 1120 | 4.497e+5 | 401.488 | 449.667 | 0.401 | 0.196 |
| MS-2 | 1186 | 4.912e+5 | 414.153 | 491.185 | 0.414 | 0.214 |
| MS-3 | 1079 | 4.210e+5 | 390.152 | 420.974 | 0.390 | 0.183 |
| SC | 1052 | 3.979e+5 | 378.261 | 397.931 | 0.378 | 0.173 |
| LA-3 | 1166 | 4.655e+5 | 399.215 | 465.485 | 0.399 | 0.202 |
| TXAR | 1835 | 7.504e+5 | 408.912 | 750.354 | 0.409 | 0.326 |
| TX | 1601 | 6.334e+5 | 395.604 | 633.362 | 0.396 | 0.275 |

ROH: Run of Homozygosity; $F_{ROH}$ : Proportion of genome covered by ROHs.

**Table 2:** Model, log likelihood, number of parameters and AIC for all *δaδi* models.

| Model | Log Likelihood | Parameters | AIC |
| --- | --- | --- | --- |
| Equil | -38193.0255956529 | 1 | 76388.0511913058 |
| Neutral | -132482.569644277 | 1 | 264967.139288554 |
| Exp growth 1D | -234.982747590169 | 2 | 473.9654951803 |
| Two epoch | -128.940800020471 | 2 | 261.8816000409 |
| Bottlegrowth 1D | -121.689448170364 | 3 | 249.3788963407 |
| Three epoch | -78.0804739920422 | 4 | 164.1609479841 |
| Three epoch inbreeding | -539.513394270092 | 5 | 1089.0267885402 |
| Four epoch | -57.3965083546937 | 6 | 126.7930167094 |
| Five epoch | -60.5216698618606 | 8 | 137.043397237 |

We summarized individual-level inbreeding using *F_ROH_*, the proportion of the genome covered by Runs of Homozygosity (ROHs), which provides a direct genomic measure of inbreeding that is robust to allele-frequency assumptions (McQuillan et al. 2008). We measured *F_ROH_* using two thresholds: >250Kb “short” ROHs and >1Mb “long” ROHs. The mean *F_ROH_*_250_ was 0.212 (*Min* = 0.137*, Max* = 0.326*, Median* = 0.205*, SD* = 0.055), indicating that approximately 21% of the average alligator genome is contained within ROHs of this size or larger. For comparison, previous studies of the critically endangered Chinese alligator report extensive genome coverage by ROHs (*F_ROH_*= 46.7%–69.6%), consistent with recent, severe demographic contraction (Yang et al. 2023).

In comparison, large ROHs (>1 Mb), which typically reflect very recent shared ancestry, comprised a much smaller fraction of the genome (*F_ROH_*_1_*_Mb_* = 0.0105)(*n* = 356*, Min* = 0.0018*, Max* = 0.0154*, Median* = 0.006*, SD* = 0.0118)(Figure 2). The combined length of large ROHs within American alligator genomes ranged from 4.190 Mb to 116.613 Mb (*n* = 356*, Mean* = 24.149*, Median* = 13.900*, SD* = 27.214). While the length of large ROHs (> 1 Mb) were highly consistent (*Mean* = 1.240*, Median* = 1.197*, SD* = 0.129), the number of segments per individual varied significantly (*Mean* = 18.737*, Median* = 12.000*, Min* = 4*, Max* = 88*, SD* = 20.150). American alligators have a much smaller *F_ROH_*_1_*_Mb_* than Chinese alligators, whose large average *F_ROH_*_1_*_Mb_* (*Mean* = 0.327*, Min* = 0.1963*, Max* = 0.5644*, SD* = 0.0661) rivals that of the critically endangered mountain gorilla (*Mean* = 0.345) (Xue et al. 2015; Pan et al. 2025).

The length class distribution of ROHs can contextualize inbreeding load within a species (Meyermans et al. 2020; Martin et al. 2023). Notably, the majority of ROHs in *A. mississippiensis* genomes fall within the short length class (250*Kb−*1*Mb*), supporting patterns of historic inbreeding instead of recent consanguineous pairings (Martin et al. 2023)(Figure 2C). Because ROH hot- and cold-spots can be produced by selection or variation in recombination landscapes, we tested for a non-random genomic distribution using a permutation test (Bosse et al. 2012; Ceballos et al. 2018; Minn et al. 2025). We did not see evidence of clustering of ROHs when comparing all ROHs (> 250Kb) (*P* = 0.456, permutation test) or long ROHs (> 1 Mb) (*P* = 0.238, permutation test)(Figure 2A).

### Demographic history inference

To infer historical changes in effective population size (*N_e_*) from genome-wide variation, we used site-frequency-spectrum (SFS)–based demographic modeling approaches that are well suited for non-model organisms. We first compared a set of alternative demographic scenarios using models implemented in the *δaδi*-cli framework, including models describing equilibrium populations, population growth, population bottlenecks, and multi-epoch demographic histories, with and without inbreeding (Gutenkunst et al. 2009; Huang et al. 2023). Because American alligators lack many of the reference resources available for classic model systems, we used *donni* to obtain biologically plausible starting parameters for each model (Tran et al. 2024). To minimize the confounding effects of selection on demographic inference, analyses were restricted to intergenic sites under the assumption that these regions are less directly affected by purifying or positive selection than coding sequence (Eyre-Walker and Keightley 2007; Boyko et al. 2008; Galtier 2016; Johri et al. 2022).

Among the default *δaδi* models, the Three-Epoch model provided the best fit to the observed SFS of intergenic sites. To refine inference of population decline dynamics, we additionally implemented custom Four-Epoch (4E) and Five-Epoch (5E) models. The Four-Epoch model fit the data significantly better than either the Three-Epoch model (*AIC*_3_*_E_* = 164.61*, AIC*_4_*_E_* = 129.793) and the Five-Epoch model (*AIC*_5_*_E_* = 137.043). The Four-Epoch model supports a demographic history characterized by a large ancestral effective population size (2.191 *×* 10^5^) followed by multiple sequential population declines to a contemporary *N_e_* of 186 (*Ne*_0_ = 2.191 *×* 10^5^*, Ne*_1_ = 1.022 *×* 10^5^*, Ne*_2_ = 3.855 *×* 10^2^*, Ne_Contemp_* = 1.866 *×* 10^2^)(Table 2).

Because different SFS-based approaches can have complementary sensitivity to demographic change across timescales, we also inferred population history using StairwayPlot2, which is particularly powerful for detecting recent demographic dynamics (Liu and Fu 2020). This method similarly supports a significant population decline since the Last Glacial Period (LGP), though estimates for median contemporary *N_e_* were much lower than those inferred using *δaδi* (*Ne_Ancient_* = 6.09*e*^6^*, Ne_LGP_* = 1.07*e*^6^*, Ne_LGM_* = 2.26*e*^5^*, Ne*_1500_*_BC_* = 4.27*e*^4^*, Ne*_1400_*_AD_* = 8.33*e*^3^*, Ne*_1900_*_AD_* = 1.58*e*^3^*, Ne_Contemp_* = 5.66*e*^1^) (Figure 3) (Hughes et al. 2013).

**Figure 3:**
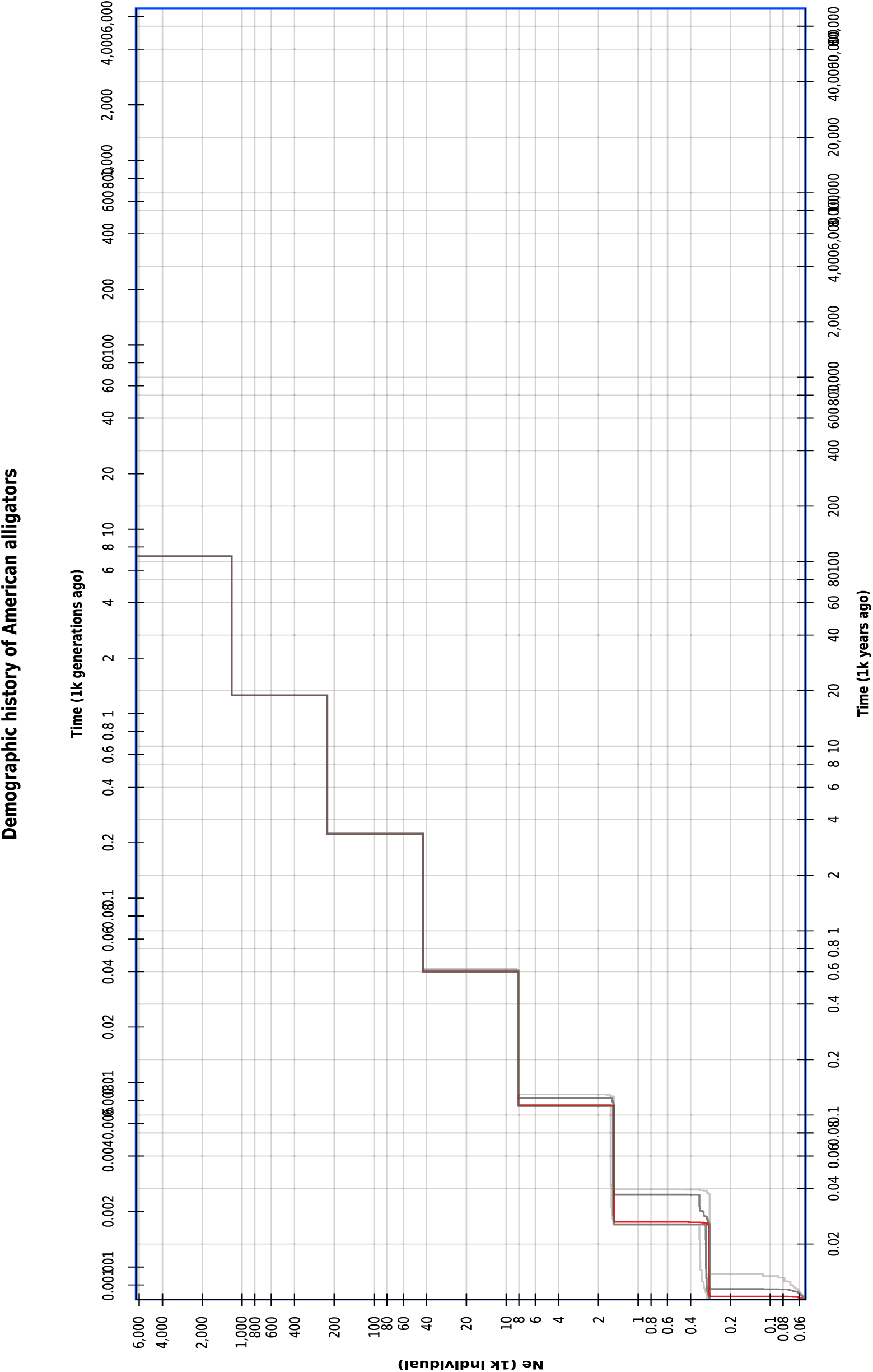
Demographic history inference via StairwayPlot2.

To ensure that individuals at the edges of alligator habitat did not have any undue influence on demographic history reconstruction, we also performed a permutation test wherein we reconstructed the demographic history excluding one individual at a time. There was no significant difference in median *N_e_* in each of the permutations, though the 97.5% and 87.5% confidence intervals were slightly broader. It is important to note that we only had data for 19 individual genomes. This may limit the resolution of *N_e_* estimates, as best practices require dozens to hundreds of samples (Beichman et al. 2017; Nadachowska-Brzyska et al. 2022). Additionally, SFS methods like those used in this study may be most accurate between the present and 30 generations before present (Patton et al. 2019). However, our patterns of population decline are also consistent between methods and with previously published demographic reconstruction of American alligators (Green et al. 2014). Thus, while ancient estimates of population size may lack precision, the overall conclusion that alligator populations were in decline before anthropogenic overhunting and habitat destruction is supported by multiple lines of evidence.

### Inference of Fine-Scale Recombination Landscape

Recombination plays a central role in shaping how genetic variation is redistributed following population bottlenecks by breaking down linkage among alleles. To characterize recombination rate variation across the genome, we used ReLERNN, a convolutional neural network-based approach that infers recombination landscapes from patterns of linkage-disequilibrium in population genomic data (Adrion et al. 2020). This analysis generated a genome-wide, sex-averaged recombination landscape in American alligators (Figure 4). The mean genome-wide recombination rate was 1.682 *×* 10*^−^*^9^ c/bp (*Median* = 1.296 *×* 10*^−^*^9^*, SD* = 1.594 *×* 10*^−^*^9^) (Figure 4). Strikingly, the distal ends of the chromosomes have much higher recombination rates than the central regions (Figure 4). This pattern persists across all 16 chromosomes, though the smallest chromosomes’ recombination landscapes are less starkly distal (Figure 4).

**Figure 4:**
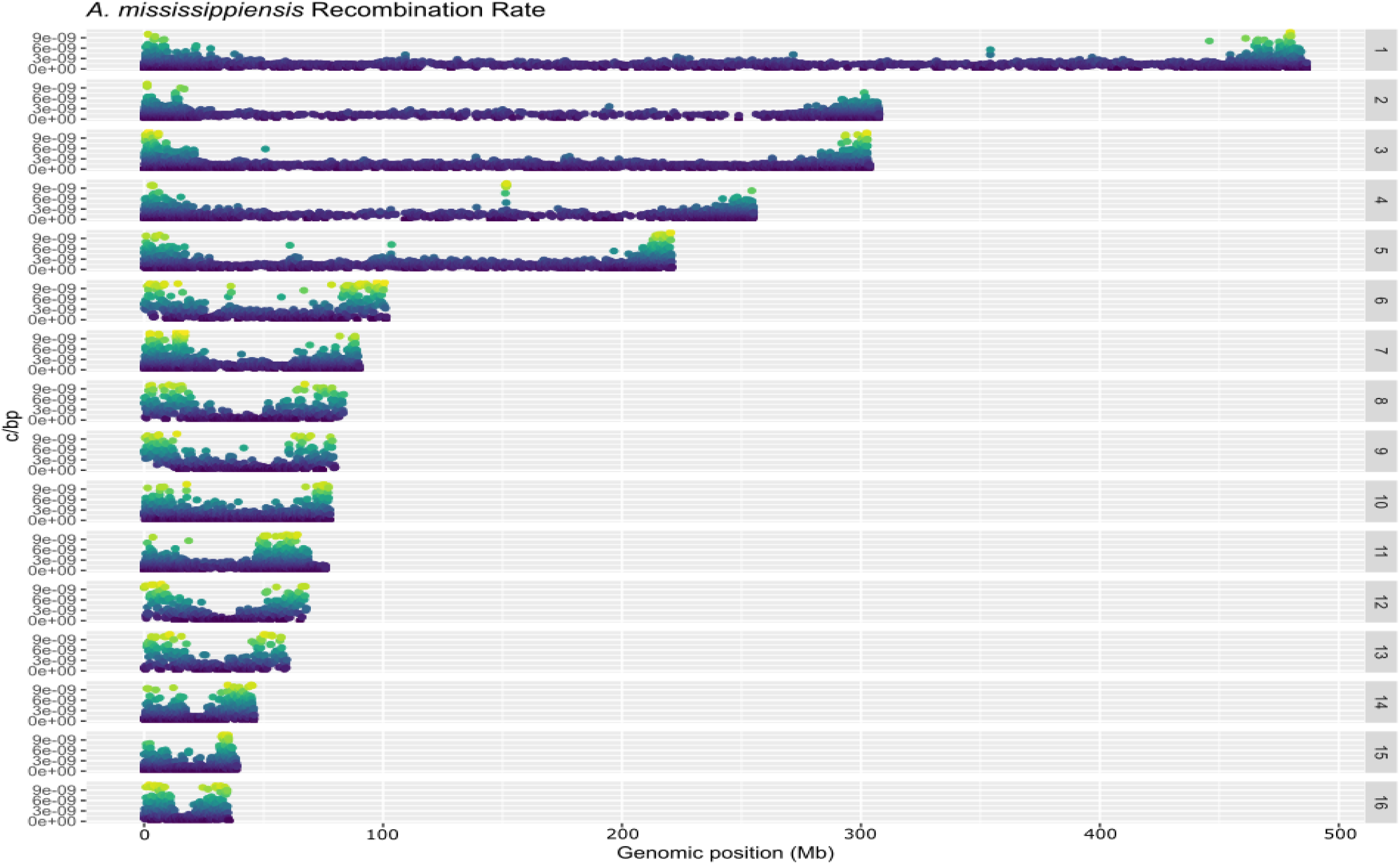
Sex-averaged recombination landscape across 16 alligator autosomes.

To evaluate whether the pronounced distal recombination bias corresponds to underlying chromosome architecture, we examined the distribution of telomeric and centromeric repeat elements in the reference genome. Using Tandem Repeats Finder (TRF) Benson (1999), we identified canonical vertebrate telomeric repeats (TTAGGG) and candidate centromeric satellite repeats (100–200 bp motifs shared across chromosomes) in the *A. mississippiensis* reference assembly (Crother et al. 2023), guided by published karyotypic information (Valleley et al. 1994). Elevated recombination rates extended far beyond the boundaries of telomeric repeat regions on all chromosomes, while recombination-suppressed regions exceeded the inferred extent of candidate centromeric repeats. These results suggest that distal-biased recombination landscape is not driven directly by telomeric or centromeric structure, though it is possible that other structural chromosome features may contribute. We note that inference of repetitive regions may be incomplete due to challenges associated with assembling repeat-rich genomic sequences.

Because recombination rate variation is often correlated with GC content, gene density, and levels of genetic diversity, we next examined relationships among these genomic features (Fullerton et al. 2001; Freudenberg et al. 2009; Tortereau et al. 2012; Mugal et al. 2013; Kiktev et al. 2018). All pairwise relationships were statistically significant but modest in magnitude. GC content was positively correlated with recombination rate, nucleotide diversity, and gene density, (*ρ_GC_*___*_REC_* = 0.150, *ρ_GC_*___*_P I_* = 0.362, *ρ_GC_*___*_GD_* = 0.403, *P <* 0.0001, Spearman correlation coefficient). Recombination rate and nucleotide diversity were also positively correlated, although the relationship was moderate (*ρ* = 0.315*, P <* 0.0001, Spearman correlation coefficient). Gene density showed only modest enrichment at the distal ends of the chromosomes, in contrast to the pronounced distal bias observed for recombination rate. Outside of these distal regions, gene-rich regions overlapped more strongly with areas of elevated GC content (*ρ* = 0.403, *P <* 0.0001, Spearman correlation coefficient). Gene density was also weakly correlated with recombination rate and nucleotide diversity (*ρ_GD_*___*_REC_* = 0.126, *ρ_GD_*___*_P I_* = 0.205, *p <* 0.0001, Spearman correlation coefficient), indicating that gene-rich regions do not strongly predict the spatial distribution of either recombination or genetic diversity. Overall, these patterns are consistent with GC-biased gene conversion (gBGC), a recombination-associated process in which mismatches during gene conversion are preferentially resolved in favor of GC alleles, leading to localized increases in GC content and generating correlations among recombination rate, nucleotide diversity, and genomic composition (Webster et al. 2006; Galtier and Duret 2007; Glémin et al. 2015; Clessin et al. 2025).

To test whether localized regions of unusually high recombination occurred outside chromosome ends, we performed a permutation analysis based on the upper 5% of genome-wide recombination rates. High-recombination windows were overwhelmingly concentrated at distal regions, limiting power to detect discrete hotspots within these areas. Using a sliding-window approach that excluded distal regions, we identified two candidate high-recombination regions (Table 4), which together contained 732 annotated genes enriched for Gene Ontology terms related to neural signaling (Table 5). Despite broad-scale correlations among recombination rate, GC content, and gene density, we find no evidence that recombination hotspots are preferentially associated with gene-rich regions. However, because recombination rates were inferred from linkage disequilibrium patterns in a relatively small sample, our ability to detect fine-scale hotspots may be limited.

## Discussion

The American alligator (*A. mississippiensis*) is widely regarded as a conservation success story, having rebounded demographically following severe population declines caused by overharvesting and habitat loss in the nineteenth and twentieth centuries. However, despite the species’ designation as “fully recovered,” larger census population sizes do not necessarily mean full recovery from a population bottleneck. Using whole genome data from across the species’ range, we show that contemporary alligator populations retain low levels of nucleotide diversity and elevated homozygosity. Interestingly, these patterns are best explained by a history of prolonged population decline rather than recent consanguineous inbreeding associated with a recent anthropogenically-driven population bottleneck and subsequent expansion.

Genome-wide measures of diversity indicate that *A. mississippiensis* harbors substantially lower standing genetic variation than most vertebrates (Chen et al. 2017; Leroy et al. 2021). Although autosomal heterozygosity estimates from whole-genome data are higher than those reported in earlier microsatellite studies (Glenn et al. 1998; Davis et al. 2002; Ryberg et al. 2002), they remain low relative to birds and other non-avian reptiles (Iannucci et al. 2021; Martin et al. 2023; Webster et al. 2024; Day et al. 2025), and only modestly exceed estimates reported for the critically endangered Chinese alligator (Pan et al. 2019; Yang et al. 2023; Pan et al. 2025). Importantly, this reduction in genetic diversity is uniformly distributed across chromosomes and individuals, indicating that it is not driven by localized genomic effects or a subset of highly inbred lineages. Together, these patterns point to long-term constraints on genetic variation rather than short-term demographic processes.

Analysis of runs of homozygosity (ROHs) provides critical temporal context for interpreting these patterns. While method-of-moments and heterozygosity-based inbreeding coefficients indicate elevated genome-wide homozygosity, ROHs in American alligators are overwhelmingly short and evenly distributed across the genome. This contrasts sharply with patterns observed in Chinese alligators and other critically endangered vertebrates, where a large proportion of the genome is contained within long ROHs (Xie et al. 2022; Stanhope et al. 2023; Hoffman et al. 2024; Lan et al. 2025). Long ROHs, which are hallmarks of recent consanguineous mating, comprise only a small fraction of American alligator genome coverage and are rare compared with those observed in species that have experienced severe, recent bottlenecks. The majority of genome coverage by ROHs (*F_ROH_*) is due to the preponderance of short ROHs, which occur as recombination breaks down long IBD segments over time. Thus, the genomic architecture of homozygosity in *A. mississippiensis* is consistent with gradual accumulation of inbreeding under long-term demographic decline rather than recent anthropogenic-driven changes.

Demographic history inference further supports this interpretation. Using complementary site-frequency-spectrum approaches, we infer a prolonged decline in effective population size of approximately three orders of magnitude (970-fold), beginning 2°1-25 kya during the late Pleistocene with continued reductions through subsequent epochs. This pattern is consistent with population contractions associated with Pleistocene climatic oscillations, which are thought to have generated latitudinal gradients in genetic diversity across many taxa (Fonseca et al. 2023). Although absolute estimates of historical and contemporary *N_e_*vary between methods — as expected given differences in sensitivity and resolution — both approaches recover concordant trajectories of long-term decline. These results align closely with previous genomic analyses of alligators and reinforce the conclusion that substantial erosion of genetic diversity likely began well before human intervention (Green et al. 2014). Furthermore, recent conservation efforts, while successful in restoring census size, may not yet have had sufficient time to generate detectable genomic recovery, particularly given the species’ long generation time of approximately 15 years (Florida Fish and Wildlife Conservation Commission 2026).

Beyond demographic history, our results highlight the importance of recombination landscape structure in shaping contemporary patterns of genetic diversity. We present the first fine-scale, genome-wide recombination map for a crocodilian and show that recombination in American alligators is strongly biased toward the distal ends of chromosomes, with large central regions experiencing reduced recombination. This pattern is consistent across all autosomes and is not directly explained by the position of telomeric or centromeric repeats, suggesting that other aspects of chromosome-scale architecture influence recombination rate variation in this species. The distal bias in recombination resembles patterns reported in some reptiles, such as snakes and lizards, but differs from the more uniform distribution of recombination observed in chickens, a sister lineage to crocodilians within Archosauria (Lisachov et al. 2017; Haenel et al. 2018; Lisachov et al. 2019; Schield et al. 2020; Weng et al. 2019).

Recombination rate heterogeneity is associated with spatial variation in nucleotide diversity, with both recombination and diversity elevated at chromosome ends. This correlation is consistent with elevated recombination promoting the maintenance of genetic diversity in distal regions by reducing the effects of linked selection, while patterns of genetic variation in the extended central regions remained shaped by drift and linkage. This architecture is expected to slow genome-wide genetic recovery following population expansion, as large portions of the genome experience limited reshuffling of alleles (Bouzat 2010; Charlesworth and Willis 2009; Kirkpatrick and Jarne 2000). Despite the role of recombination in breaking up long ROHs, the distribution of ROHs shows little evidence of strong spatial structuring across the genome. Notably, while recombination rate, GC content, gene density, and nucleotide diversity are moderately correlated at broad scales, we find no evidence for discrete recombination hotspots associated with gene-rich regions, distinguishing the alligator recombination landscape from that of birds and mammals (Fullerton et al. 2001; Galtier et al. 2001; Galtier 2004; Duret and Galtier 2009; Auton et al. 2013; Tortereau et al. 2012; Singhal et al. 2015).

There is evidence that crocodilians may exhibit extreme sex differences in recombination rate (heterochiasmy), as the linkage map of female saltwater crocodiles is approximately sixfold larger than that of males (Miles et al. 2009). This female-to-male recombination ratio rivals that observed in fish and amphibians, but greatly exceeds that seen in birds, the closest living relatives of crocodilians (Campos-Ramos et al. 2009; Hansson et al. 2010; Malinovskaya et al. 2020; Tan et al. 2024; Xie et al. 2025). Importantly, sex differences in recombination rate are driven by cellular mechanisms following sex determination, so the absence of genetic sex determination in this clade does not explain the disparity in map size (Lynn et al. 2002; Tease and Hultén 2004; Lynn et al. 2005). Because our recombination map is sex-averaged, it remains unclear whether sex-specific recombination differences contribute to the observed distal bias in American alligators. Resolving this question will require pedigreed samples or sex-specific linkage data.

Together, our results highlight how the interaction between long-term demographic history and chromosome-scale recombination structure shapes genome-wide patterns of diversity, with important implications for understanding genetic recovery in species of conservation concern. Although population numbers have rebounded dramatically, genomic signatures of past bottlenecks remain widespread, and recombination heterogeneity likely constrains the pace and spatial distribution of genetic recovery. These results emphasize that demographic recovery alone does not guarantee restoration of genetic diversity and underscore the importance of incorporating genome-wide measures of genetic health into conservation assessments. More broadly, our study illustrates how interactions between demographic history and genome architecture can shape evolutionary potential long after populations appear demographically “recovered.”

## 1 Methods

### 1.1 Molecular methods

#### DNA collection and sequencing

We received previously collected tissue samples from 19 alligators from throughout their range (Supplementary Table 1, Supplementary Figure 2). Because these samples were collected from alligators that had previously been hunted or euthanized due to human-alligator interactions, no IACUC approval was necessary.

We extracted genomic DNA from tissue and blood samples using a Qiagen DNeasy Blood and Tissue Kit (Qiagen, Cat. No. 69504). Only samples with a minimum genomic DNA concentration of 12.5 *ng*/*µL*, *≥* 2*µg* total DNA, and a minimum volume of 15 *µL* were sent for sequencing.

Whole genome sequencing and library construction was done by Innomics (IDSEQ Inc., Sunnyvale, CA). Briefly, sequencing was done using the DNBseq platform, with a read length of PE150. Raw reads were filtered, removing adaptor sequences, contamination, and low-quality reads. Samples were sequenced at 30x coverage on one lane of an Illumina Novaseq S4 Flow Cell.

#### Read Mapping & Variant calling

We followed GATK best practices for data pre-processing and germline variant discovery as previously described (DePristo et al. 2011; Jensen et al. 2021). Briefly, we created a sequence dictionary for the alligator reference genome using the CreateSequenceDictionary command. We then mapped our reads against the genome using bwa mem Li (2013) and sorted the resulting SAM files using the SortSam command from Picard (Pic 2018). Using PICARD again, we identified duplicate reads before sorting and indexing them using SAMtools (Li et al. 2009). We used the HaplotypeCaller function from GATK to phase haplotypes and create an intermediate GVCF file for each sample. We then assembled a GenomicsDB datastore combining GVCF files for each sample and used this datastore for joint-calling of SNPs using the GenotypeGVCFs function. Importantly, we were unable to perform base quality score recalibration or variant quality score recalibration, due to a lack of known variant files for alligators. We used the VariantFiltration command to apply GATK’s recommended hard-filtering parameters (QD *<* 2.0, QUAL *<* 30.0, SOR *>* 3.0 FS *>* 60.0, MQ *<* 40.0, MQRankSum *<* −12.5, ReadPosRankSum *<* −8.0) to our final VCF file. We used BCFtools to generate IDs for each SNP, using the annotate command and –set-id flag.

#### Kinship

We used King to calculate relatedness between individual alligators using a PLINK binary dataset (.bim, .fam, and bed files)(Purcell et al. 2007; Manichaikul et al. 2010). We set the maximum identifiable relationship to 4th degree relatives following this command: king -b alligator19.bed –related –degree 4.

### 1.2 Population genetics methods

We used autosomal heterozygosity as a measure for genetic diversity. Measuring heterozygosity through SNPs alone can be biased by low sample size numbers, which is an important limitation of this study (Schmidt et al. 2021). Instead, we followed the recommendations of Schmidt et al. (2021), calculating heterozygosity at monomorphic and polymorphic sites. We also calculated SNP density (SNP/kb) across the population to estimate variance within the population by dividing number of SNPs by the assembled genome length (2.1 Gb). (Jensen et al. 2021).

#### Genetic diversity

In order to measure whole genome heterozygosity, we generated FASTA files from our intermediate sorted and indexed BAM files, which had duplicates marked. We then took these FASTA files and used Jellyfish 2.2.10 to generate a distribution of k-mers in each sequence (Marçais and Kingsford 2011). We used a k-mer length of 21, 100Mb RAM, and 16 threads for each FASTA file. After acquiring the .histo files, we used GenomeScope to estimate genome size from the reads, read error rate, and whole-genome heterozygosity (Ranallo-Benavidez et al. 2020). Comparing genome size estimated from GenomeScope against the reference assembly acted as an additional quality control step to identify genomes that had exceptionally high repeats or truncation.

While it is important to have a measure of whole-genome heterozygosity, knowing the genetic diversity of variable sites within a population is crucial to understanding patterns of genetic variation within the genome. We used the PLINK –het flag to calculate expected and observed heterozygosity (Chang et al. 2015).

We used VCFTools to calculate nucleotide diversity *π* along the genome in 5 MB windows using the –window-pi flag (Danecek et al. 2011). This function returns average nucleotide diversity within the windows by inferring monomorphic sites from number of SNPs/length of window.

#### Inbreeding

We calculated two inbreeding coefficients: Wright’s inbreeding coefficient, F_IT_ (Wright 1922) and the genomic inbreeding coefficient, F_ROH_ (McQuillan et al. 2008). F_ROH_ was calculated as *F_ROH_*= Σ *L_ROH_*/*L_T otal_* for both long and short ROH classes (McQuillan et al. 2008). We used both VCFTools and PLINK 1.9 to calculate *F_IT_* and the method of moments inbreeding coefficient (Danecek et al. 2011; Chang et al. 2015; Purcell et al. 2007).

#### ROH identification and analysis

Long runs of homozygosity (ROHs) indicate the presence of recent inbreeding (Gibson et al. 2006). However, shorter ROHs can indicate the presence of a historic genetic bottleneck (Martin et al. 2023). Following suggestions by Meyermans et al. (2020), we calculated SNP density for ROH analysis using VCFTools and the –SNPDensity 1000 flag, giving us a value of 2.257 SNP/kb, or 0.443 kb/SNP (Meyermans et al. 2020; Danecek et al. 2011). In order to investigate both recent and historic inbreeding, we performed two ROH analyses: one for detecting large ROHs (*>* 1 Mb), and one for detecting smaller, more ancient ROHs (*>* 250 kb), as done in (Martin et al. 2023). We used the following parameters in the –homozyg function of PLINK: –homozyg-window-snp 50, –homozyg-density 50, –homozyg-gap 200, –homozyg-window-het 2, –homozyg-window-missing 5. We set –homozyg-kb to 250 for the short ROH analysis and 1000 for the large ROH analysis.

To determine the distribution of ROHs, we calculated length classes by modifying the R code previously published for pipits (0–0.25Mb, 0.25-0.5Mb, 0.5-1.0Mb, 1.0-2.0Mb, 2.0-4.0 Mb, and > 4.0 Mb) (Martin et al. 2023). There were no ROHs larger than 4Mb. Briefly, we calculated TMRCA as L = 100/(2 t), where t is the time since the most recent common ancestor in generations. For our alligators, this equates to 250Kb = <200 generations ago, 500kb = 100 generations ago, 1Mb = 50 gens, 2 Mb = 25 gens, 4Mb = 1°2.5 gens.

#### Demographic history models

We used *δaδi*-cli to perform demographic history simulations using the following models: bottlegrowth_1d, growth, snm_1d, three_epoch, three_epoch_inbreeding, two_epoch, and equil (Huang et al. 2023). We followed the demographic inference tutorial provided by the *δaδi*-cli documentation using a folded allele frequency spectrum. We operated under the assumption that our genomes come from one population, as we do not have enough individuals to do more regional-specific inference. In order to obtain reasonable starting parameters for the *δaδi* models, we used *donni*, which is a supervised machine learning model trained by *δaδi*-simulated data (Gutenkunst et al. 2009; Tran et al. 2024). This allows for quick predictions of model parameters, which can then be plugged into *δaδi* as a reasonable start point for demographic inference. We also used the *donni* suite of tools to train our custom 4- and 5- epoch models before applying them to our data. Once we had starting parameters, we began with a global parameter fit using 25% of perturbations for each model and set our convergence criteria to 0.003.

StairwayPlot2 (v2.1.2) is an effective method for both ancient and recent population size inference (Liu and Fu 2020). We first used the *δaδi* GenerateFS function to retrieve the folded site frequency spectrum for our genomes, since we do not know the ancestral alleles (Huang et al. 2023; Gutenkunst et al. 2009). Because StairwayPlot2 requires a mutation rate and generation time, we used previous data from Green et al. (2014), (*µ* = 7.9*e −* 9, g = 15). We used the default folded SFS blueprint file, substituting the following parameters: nseq = 38, L = 5688779, random seed = 42, mu = 7.9e-9, year_per_generation = 15, xrange = 0.01, 10000.

#### Recombination landscape

ReLERNN uses recurrent neural networks to return a sex-averaged recombination landscape (Adrion et al. 2020). ReLERNN is effective in inferring recombination data from small numbers of individual genomes and is robust to missing data and incorrect demographic model specification (Adrion et al. 2020). We used our VCF containing SNPs from all 16 autosomes anad all 19 individuals. We did not have phased genomes, and thus used the –unphased flag. We used all default settings and performed a bias correction on the output recombination landscapes to account for network bias in predicting model parameters (Adrion et al. 2020).

Because of the distal pattern of the recombination landscape, we wanted to determine whether recombination was constrained in some way around telomeric or centromeric regions. We used the canonical vertebrate telomeric repeat “TTAGGG” to identify telomeric regions. To find potential centromeric regions, we first split the reference genome into scaffolds, keeping those corresponding to chromosomes 1-16. We then used Tandem Repeats Finder (TRF) (Benson 1999) to identify potential repetitive elements within each chromosome and filtered for the following criteria: motif lengths of 120-180bp, greater than 20 copies, and block lengths greater than 3kb. It is important to note that these are rough estimates of where these features may be located, and require more vigorous investigation.

#### GC Content and Gene Richness

We measured the GC content across the genome using the Bioconductor libraries GenomicRanges, GenomicDistributions (Huber et al. 2015). We also measured gene richness across the genome. Briefly, we used a sliding window approach to count the number of genes within each 100Kb window, then plotted that incidence against genomic position.

## Statistical analysis

All statistical analysis was performed in R 4.5.1 (Posit team 2025). We used the following R packages for handling genomic data: GenomicRanges, regioneR, BSgenomeForge, Biostrings, GenomicDistributions, BSgenome, windowscanr; for statistical analysis: tidyverse, lme4, broom, forcats; for plotting/graphs: gg-plot2, RColorBrewer, viridis, scales, ggridges, gghalves; for mapping: ggmap, maps, mapdata (Lawrence et al. 2013; Gel et al. 2016; Pagès 2017; H. Pagès 2017; Pagès et al. 2025; Kupkova et al. 2022; Tavares 2025; Wickham et al. 2019; Wickham and Software 2025; Wickham et al. 2025; Bates et al. 2025; Wilke 2025; Kahle et al. 2025; Brownrigg 2022; Tiedemann 2022).

## Acknowledgments

We thank Jean-Francois Gout, Federico Hoffmann, Mark Welch, and members of the Dapper and Gout labs for constructive feedback on study design and interpretation of results. We thank Marilyn Mason and members of the Hoffman and Parrott labs for aid in receiving tissue samples. This work was supported by NSF CAREER 2143063 to ALD, Mississippi State University Strategic Research Initiative seed funding to ALD.

## Data Accessibility and Benefit-Sharing

Whole genome sequencing data is available from NCBI Sequence Read Archive (SRA) at accession numbers: Data analysis and scripts used can be accessed from Dryad at the following repository:

## Author Contributions

TSG lead data collection, data analysis and writing. TSG and ALD collaborated on study design, data analysis, and writing. JK, EH, and BP contributed to sample collection and study design.

---

**Supplementary Figure 1:**
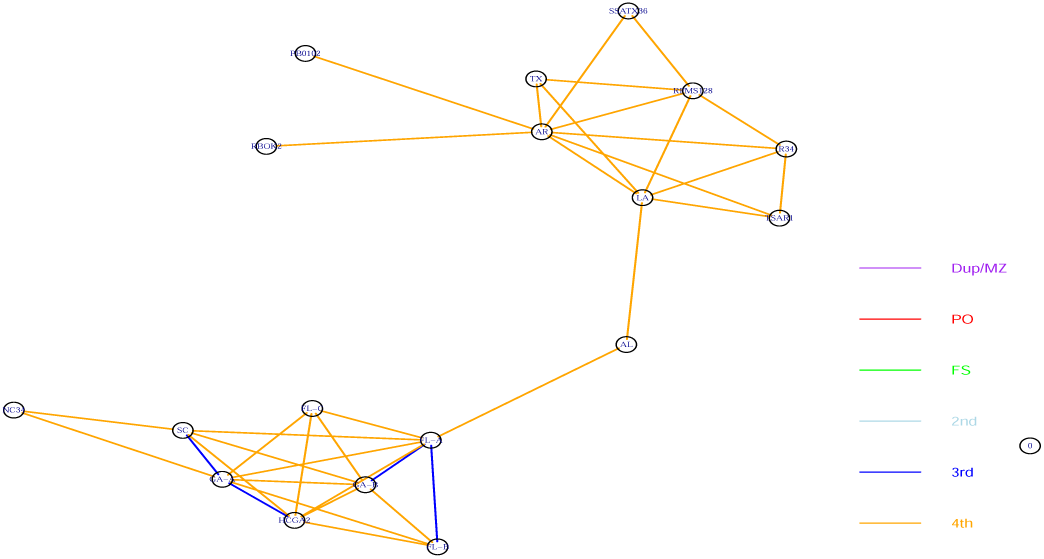
Relatedness between individuals. Orange lines represent fourth degree relation-ships, while blue lines represent third degree relationships.

**Supplementary Figure 2:**
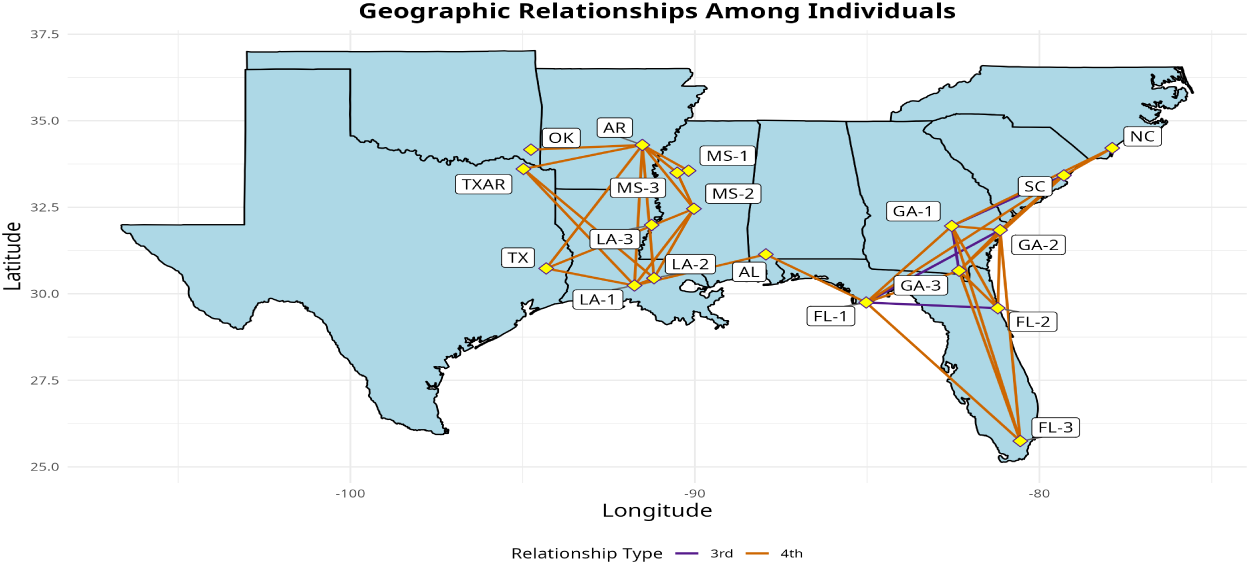
Relatedness between individuals, plotted against geographic range. Orange lines represent fourth degree relationships, while blue lines represent third degree relationships.

## Supplementary Tables

**Supplementary Table 1:**
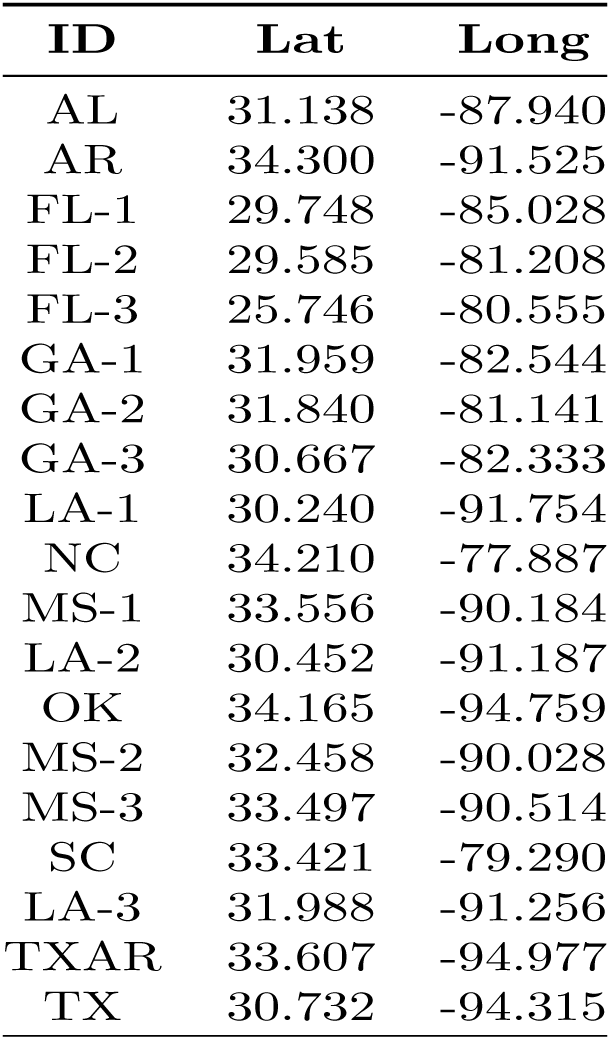
Latitude and longitude for all sampled individuals.

| ID | Lat | Long |
| --- | --- | --- |
| AL | 31.138 | -87.940 |
| AR | 34.300 | -91.525 |
| FL-1 | 29.748 | -85.028 |
| FL-2 | 29.585 | -81.208 |
| FL-3 | 25.746 | -80.555 |
| GA-1 | 31.959 | -82.544 |
| GA-2 | 31.840 | -81.141 |
| GA-3 | 30.667 | -82.333 |
| LA-1 | 30.240 | -91.754 |
| NC | 34.210 | -77.887 |
| MS-1 | 33.556 | -90.184 |
| LA-2 | 30.452 | -91.187 |
| OK | 34.165 | -94.759 |
| MS-2 | 32.458 | -90.028 |
| MS-3 | 33.497 | -90.514 |
| SC | 33.421 | -79.290 |
| LA-3 | 31.988 | -91.256 |
| TXAR | 33.607 | -94.977 |
| TX | 30.732 | -94.315 |

**Supplementary Table 2:**
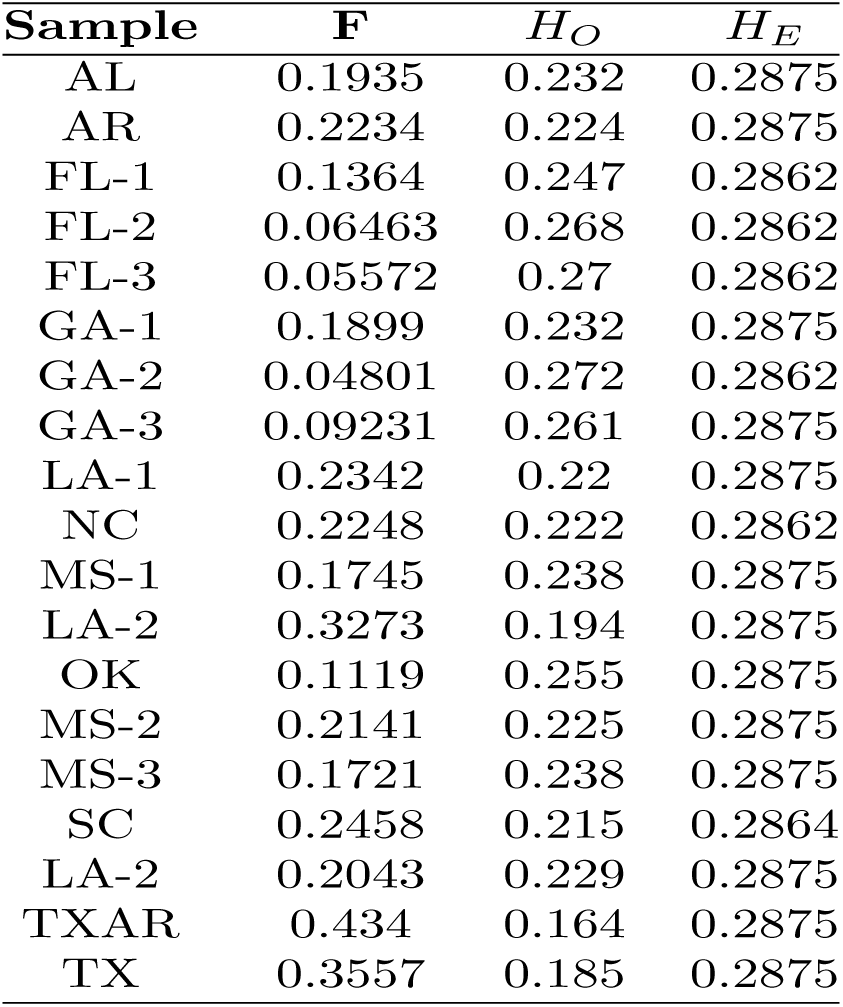
Inbreeding coefficients, expected heterozygosity, and observed heterozygosity in *A. mississippiensis* genomes.

**Supplementary Table 3:**
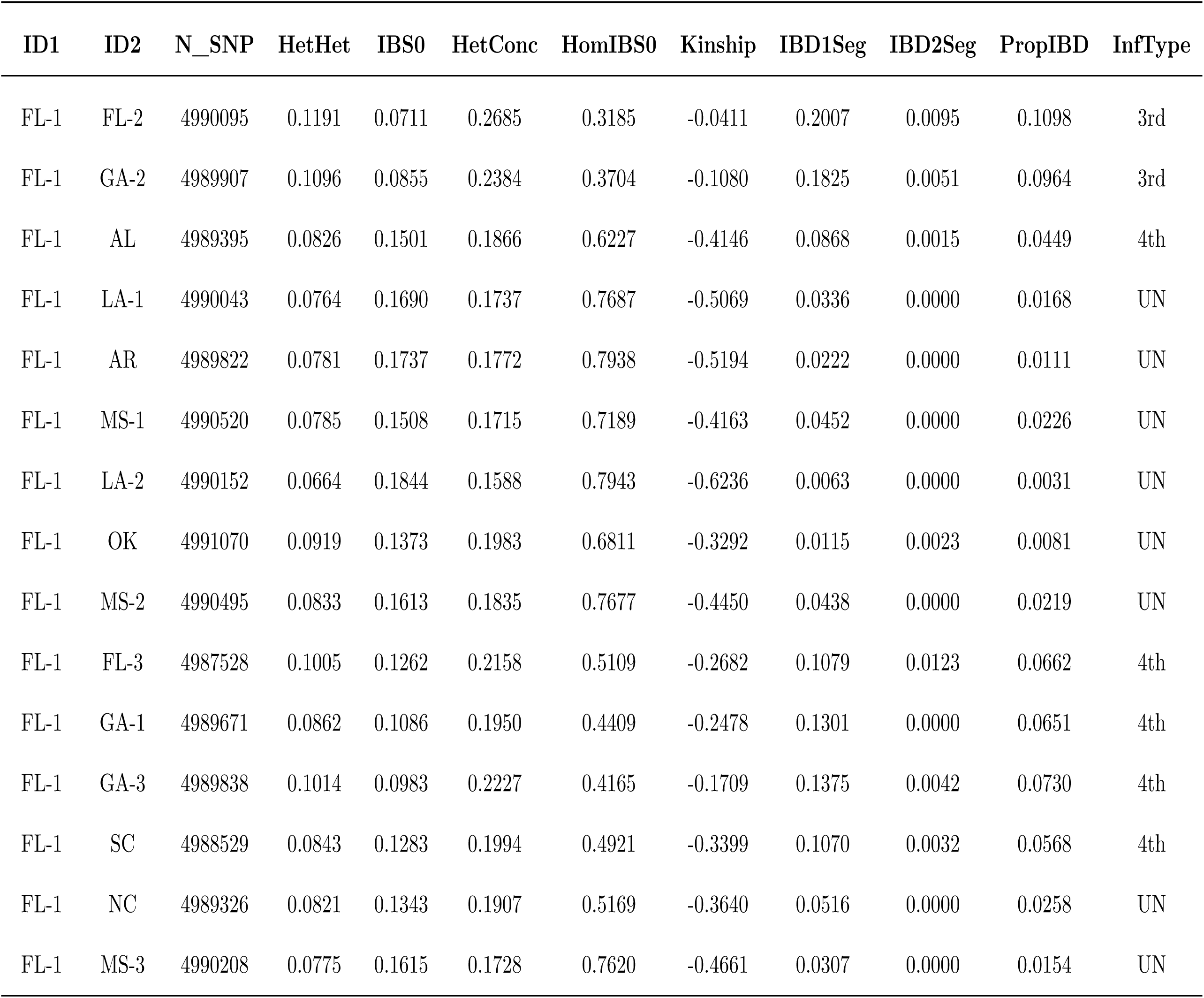
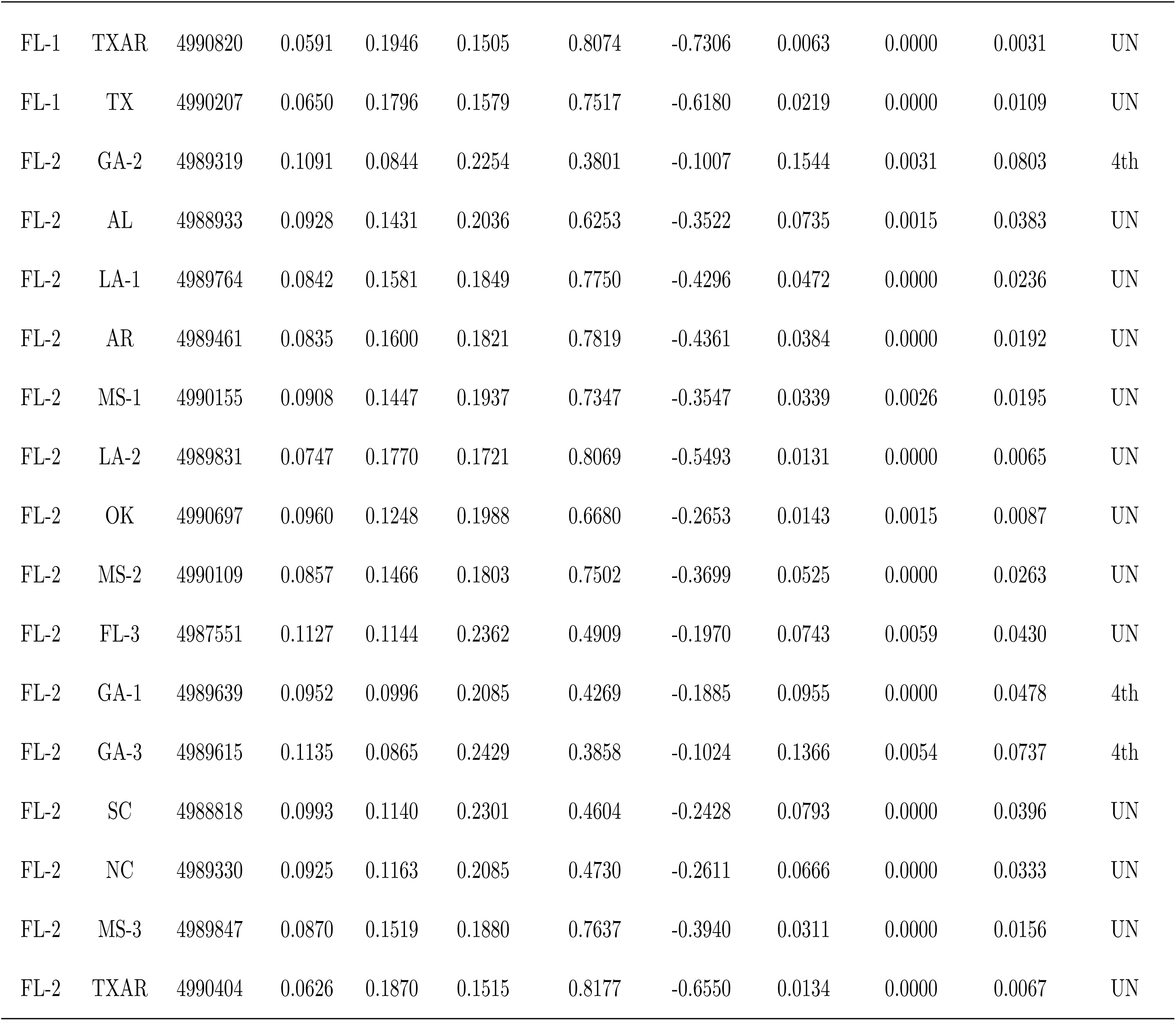
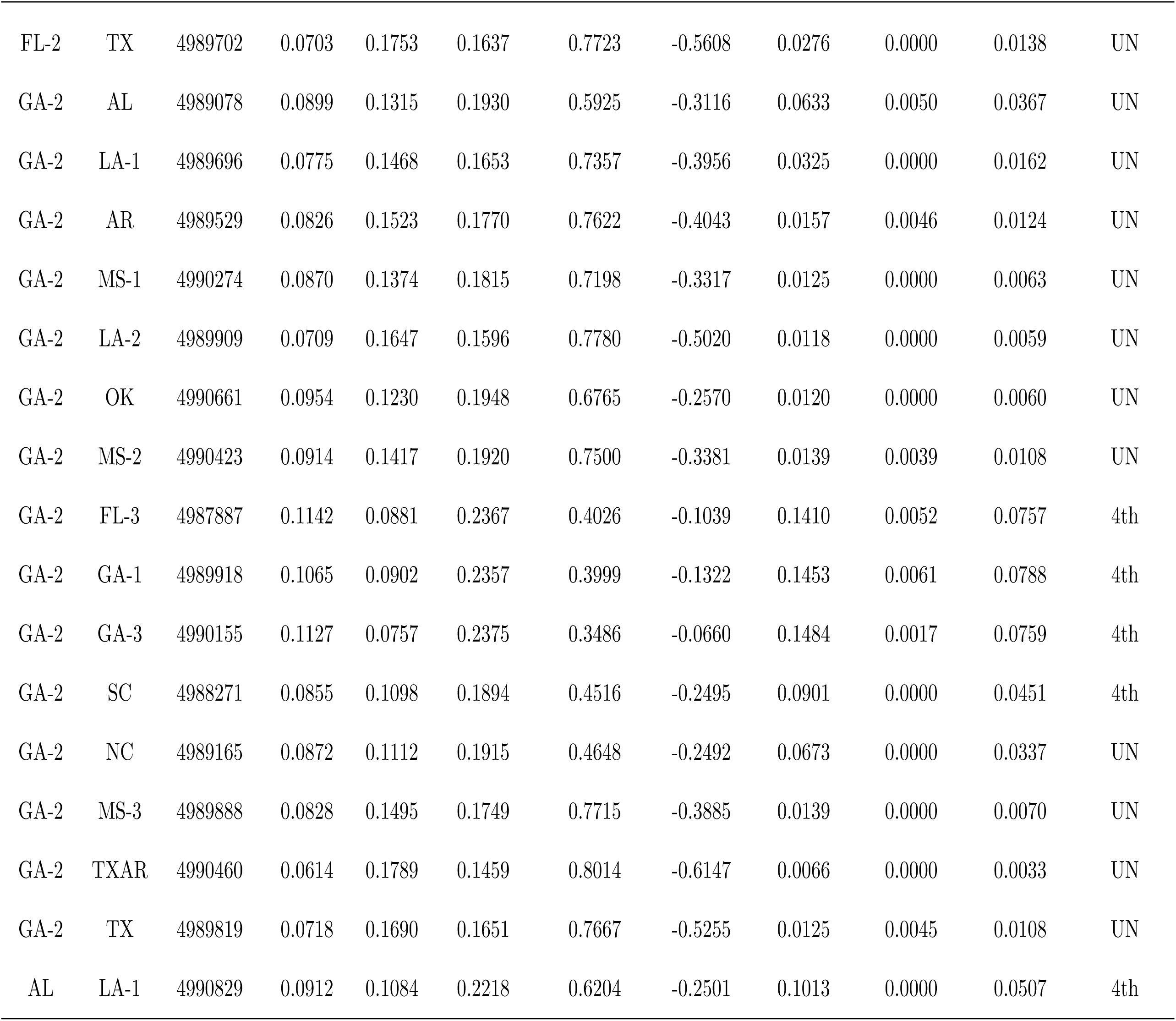
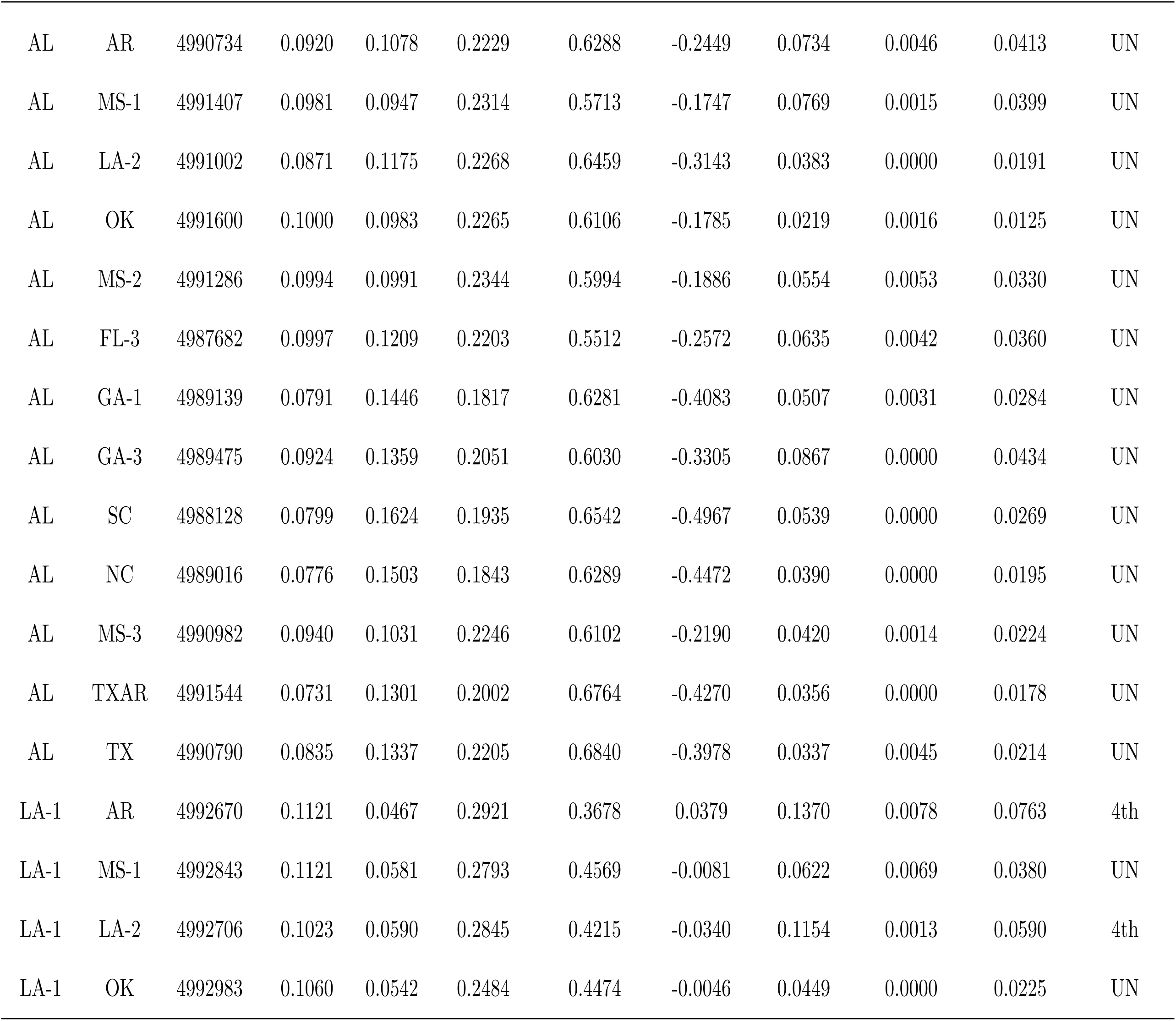
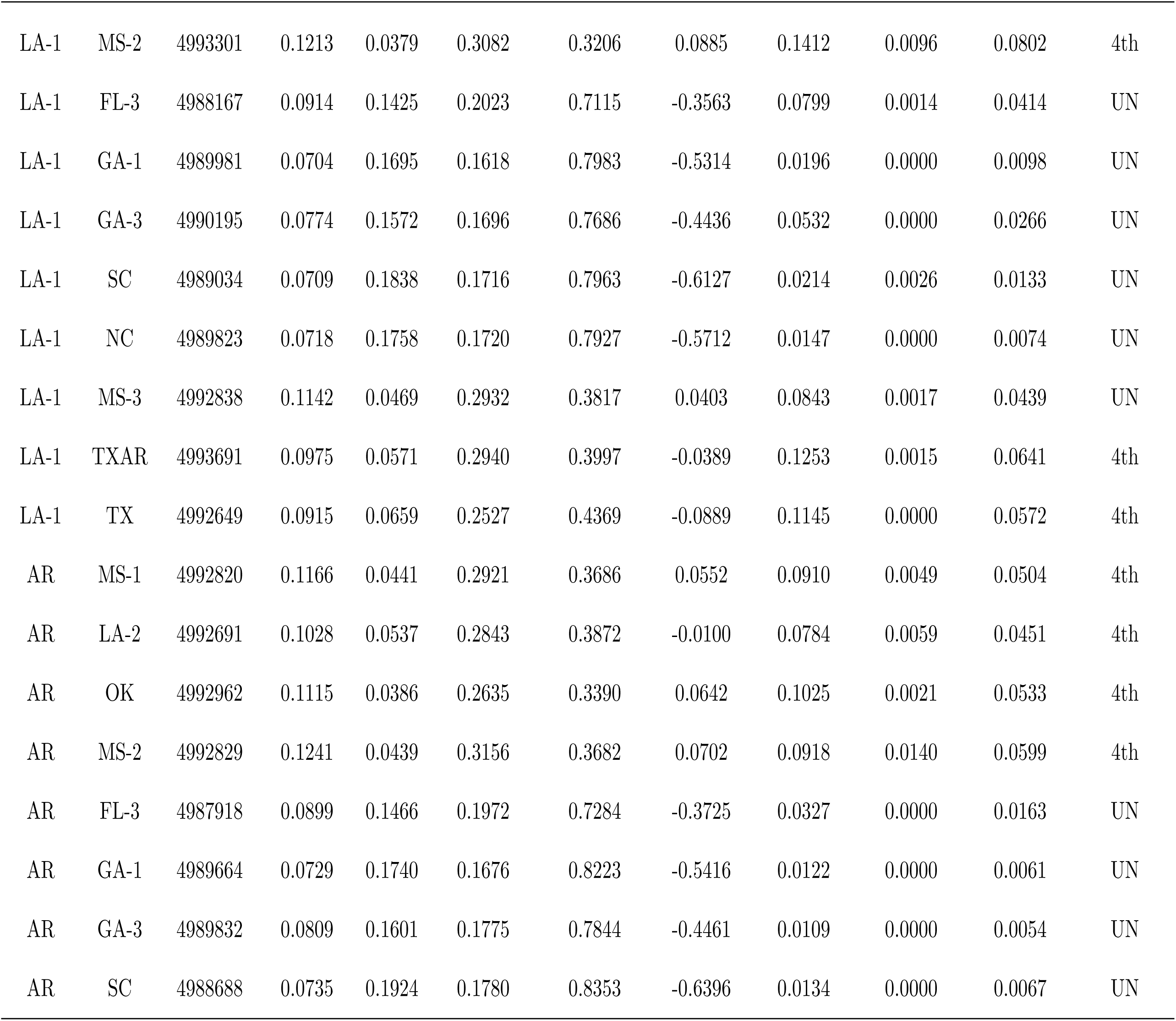
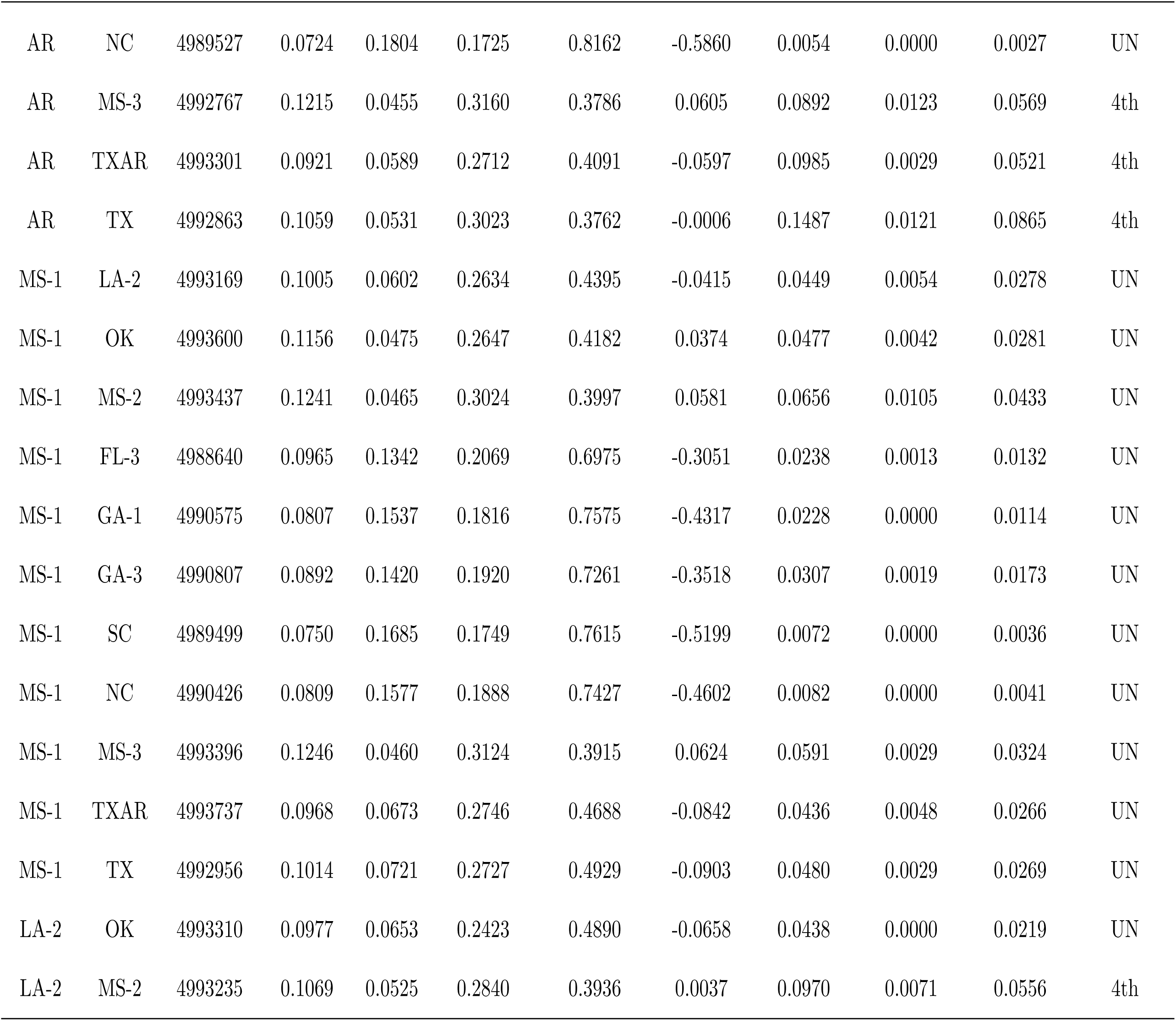
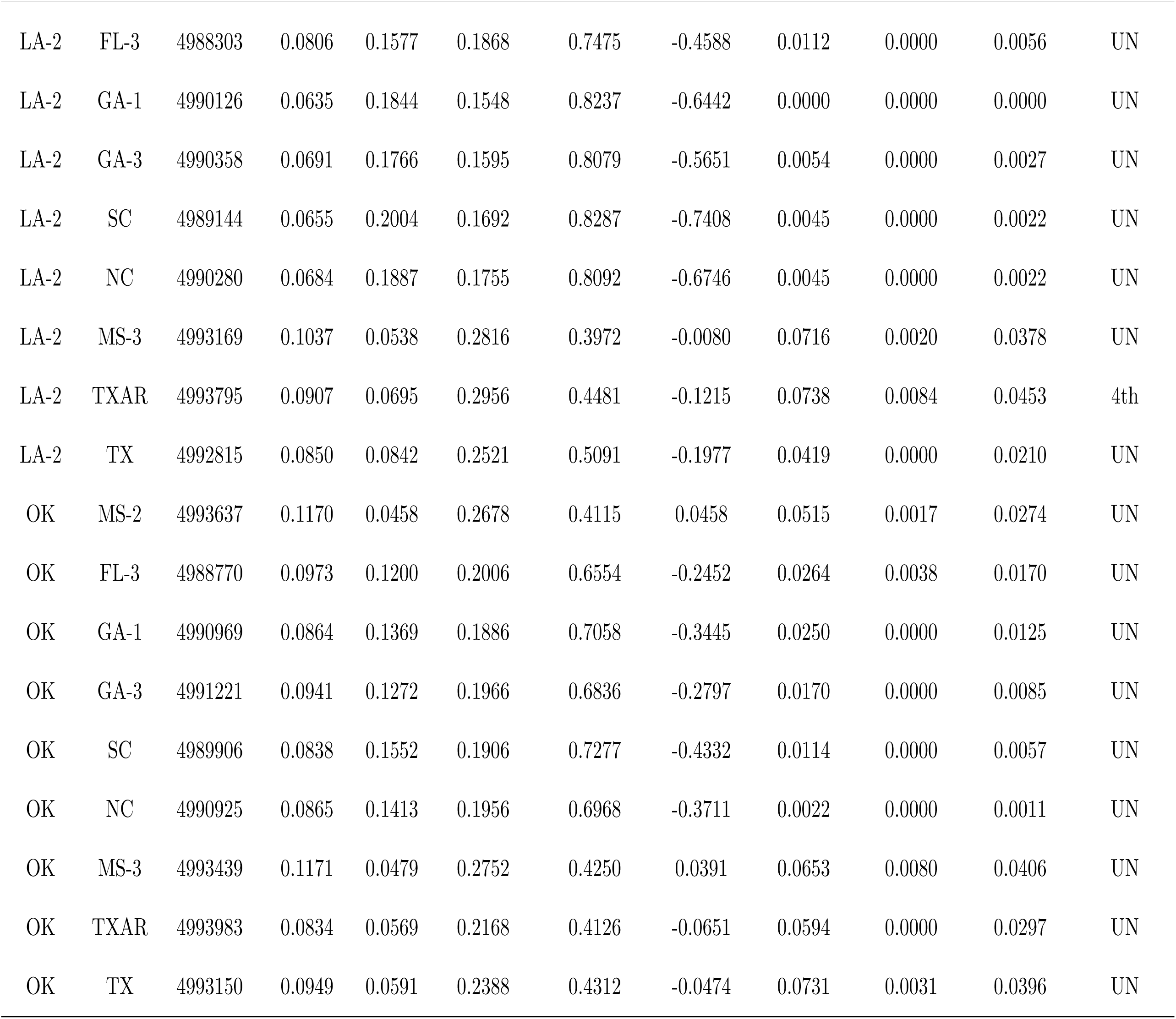
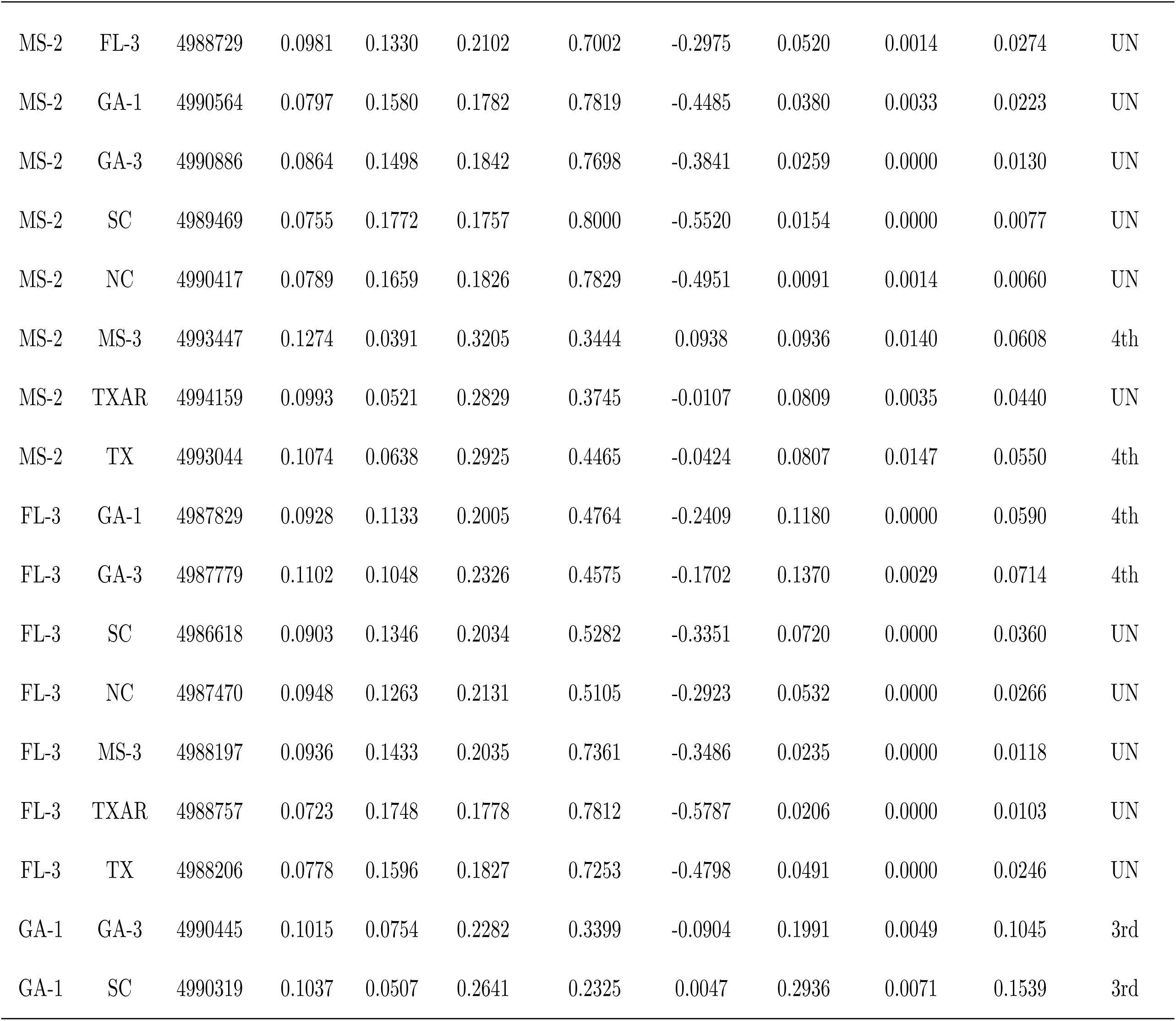
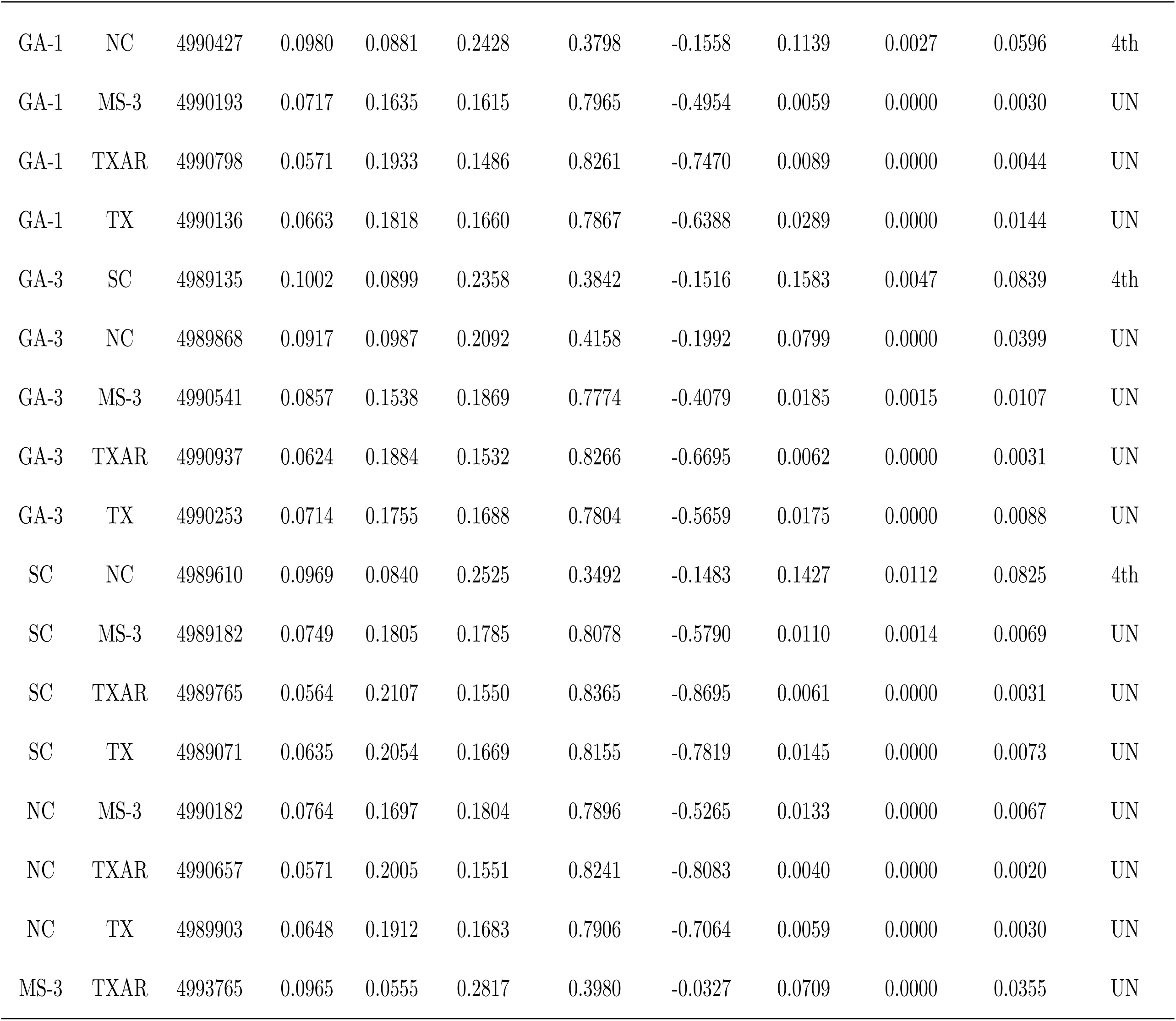
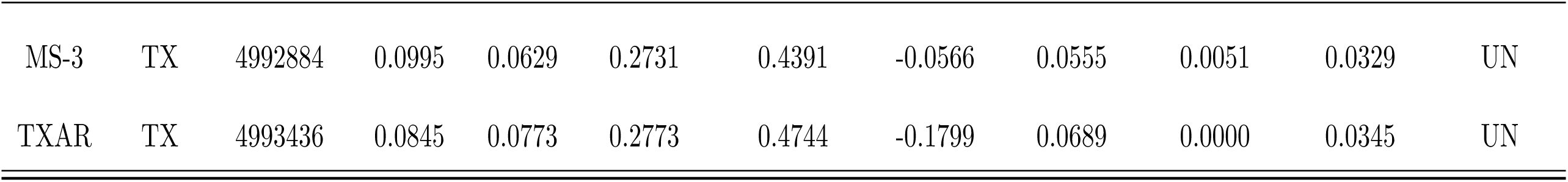
KING pairwise kinship coefficients for individual genomes.

| ID1 | ID2 | N_SNP | HetHet | IBS0 | HetConc | HomIBS0 | Kinship | IBD1Seg | IBD2Seg | PropIBD | InfType |
| --- | --- | --- | --- | --- | --- | --- | --- | --- | --- | --- | --- |
| FL-1 | FL-2 | 4990095 | 0.1191 | 0.0711 | 0.2685 | 0.3185 | -0.0411 | 0.2007 | 0.0095 | 0.1098 | 3rd |
| FL-1 | GA-2 | 4989907 | 0.1096 | 0.0855 | 0.2384 | 0.3704 | -0.1080 | 0.1825 | 0.0051 | 0.0964 | 3rd |
| FL-1 | AL | 4989395 | 0.0826 | 0.1501 | 0.1866 | 0.6227 | -0.4146 | 0.0868 | 0.0015 | 0.0449 | 4th |
| FL-1 | LA-1 | 4990043 | 0.0764 | 0.1690 | 0.1737 | 0.7687 | -0.5069 | 0.0336 | 0.0000 | 0.0168 | UN |
| FL-1 | AR | 4989822 | 0.0781 | 0.1737 | 0.1772 | 0.7938 | -0.5194 | 0.0222 | 0.0000 | 0.0111 | UN |
| FL-1 | MS-1 | 4990520 | 0.0785 | 0.1508 | 0.1715 | 0.7189 | -0.4163 | 0.0452 | 0.0000 | 0.0226 | UN |
| FL-1 | LA-2 | 4990152 | 0.0664 | 0.1844 | 0.1588 | 0.7943 | -0.6236 | 0.0063 | 0.0000 | 0.0031 | UN |
| FL-1 | OK | 4991070 | 0.0919 | 0.1373 | 0.1983 | 0.6811 | -0.3292 | 0.0115 | 0.0023 | 0.0081 | UN |
| FL-1 | MS-2 | 4990495 | 0.0833 | 0.1613 | 0.1835 | 0.7677 | -0.4450 | 0.0438 | 0.0000 | 0.0219 | UN |
| FL-1 | FL-3 | 4987528 | 0.1005 | 0.1262 | 0.2158 | 0.5109 | -0.2682 | 0.1079 | 0.0123 | 0.0662 | 4th |
| FL-1 | GA-1 | 4989671 | 0.0862 | 0.1086 | 0.1950 | 0.4409 | -0.2478 | 0.1301 | 0.0000 | 0.0651 | 4th |
| FL-1 | GA-3 | 4989838 | 0.1014 | 0.0983 | 0.2227 | 0.4165 | -0.1709 | 0.1375 | 0.0042 | 0.0730 | 4th |
| FL-1 | SC | 4988529 | 0.0843 | 0.1283 | 0.1994 | 0.4921 | -0.3399 | 0.1070 | 0.0032 | 0.0568 | 4th |
| FL-1 | NC | 4989326 | 0.0821 | 0.1343 | 0.1907 | 0.5169 | -0.3640 | 0.0516 | 0.0000 | 0.0258 | UN |
| FL-1 | MS-3 | 4990208 | 0.0775 | 0.1615 | 0.1728 | 0.7620 | -0.4661 | 0.0307 | 0.0000 | 0.0154 | UN |
| FL-1 | TXAR | 4990820 | 0.0591 | 0.1946 | 0.1505 | 0.8074 | -0.7306 | 0.0063 | 0.0000 | 0.0031 | UN |
| FL-1 | TX | 4990207 | 0.0650 | 0.1796 | 0.1579 | 0.7517 | -0.6180 | 0.0219 | 0.0000 | 0.0109 | UN |
| FL-2 | GA-2 | 4989319 | 0.1091 | 0.0844 | 0.2254 | 0.3801 | -0.1007 | 0.1544 | 0.0031 | 0.0803 | 4th |
| FL-2 | AL | 4988933 | 0.0928 | 0.1431 | 0.2036 | 0.6253 | -0.3522 | 0.0735 | 0.0015 | 0.0383 | UN |
| FL-2 | LA-1 | 4989764 | 0.0842 | 0.1581 | 0.1849 | 0.7750 | -0.4296 | 0.0472 | 0.0000 | 0.0236 | UN |
| FL-2 | AR | 4989461 | 0.0835 | 0.1600 | 0.1821 | 0.7819 | -0.4361 | 0.0384 | 0.0000 | 0.0192 | UN |
| FL-2 | MS-1 | 4990155 | 0.0908 | 0.1447 | 0.1937 | 0.7347 | -0.3547 | 0.0339 | 0.0026 | 0.0195 | UN |
| FL-2 | LA-2 | 4989831 | 0.0747 | 0.1770 | 0.1721 | 0.8069 | -0.5493 | 0.0131 | 0.0000 | 0.0065 | UN |
| FL-2 | OK | 4990697 | 0.0960 | 0.1248 | 0.1988 | 0.6680 | -0.2653 | 0.0143 | 0.0015 | 0.0087 | UN |
| FL-2 | MS-2 | 4990109 | 0.0857 | 0.1466 | 0.1803 | 0.7502 | -0.3699 | 0.0525 | 0.0000 | 0.0263 | UN |
| FL-2 | FL-3 | 4987551 | 0.1127 | 0.1144 | 0.2362 | 0.4909 | -0.1970 | 0.0743 | 0.0059 | 0.0430 | UN |
| FL-2 | GA-1 | 4989639 | 0.0952 | 0.0996 | 0.2085 | 0.4269 | -0.1885 | 0.0955 | 0.0000 | 0.0478 | 4th |
| FL-2 | GA-3 | 4989615 | 0.1135 | 0.0865 | 0.2429 | 0.3858 | -0.1024 | 0.1366 | 0.0054 | 0.0737 | 4th |
| FL-2 | SC | 4988818 | 0.0993 | 0.1140 | 0.2301 | 0.4604 | -0.2428 | 0.0793 | 0.0000 | 0.0396 | UN |
| FL-2 | NC | 4989330 | 0.0925 | 0.1163 | 0.2085 | 0.4730 | -0.2611 | 0.0666 | 0.0000 | 0.0333 | UN |
| FL-2 | MS-3 | 4989847 | 0.0870 | 0.1519 | 0.1880 | 0.7637 | -0.3940 | 0.0311 | 0.0000 | 0.0156 | UN |
| FL-2 | TXAR | 4990404 | 0.0626 | 0.1870 | 0.1515 | 0.8177 | -0.6550 | 0.0134 | 0.0000 | 0.0067 | UN |
| FL-2 | TX | 4989702 | 0.0703 | 0.1753 | 0.1637 | 0.7723 | -0.5608 | 0.0276 | 0.0000 | 0.0138 | UN |
| GA-2 | AL | 4989078 | 0.0899 | 0.1315 | 0.1930 | 0.5925 | -0.3116 | 0.0633 | 0.0050 | 0.0367 | UN |
| GA-2 | LA-1 | 4989696 | 0.0775 | 0.1468 | 0.1653 | 0.7357 | -0.3956 | 0.0325 | 0.0000 | 0.0162 | UN |
| GA-2 | AR | 4989529 | 0.0826 | 0.1523 | 0.1770 | 0.7622 | -0.4043 | 0.0157 | 0.0046 | 0.0124 | UN |
| GA-2 | MS-1 | 4990274 | 0.0870 | 0.1374 | 0.1815 | 0.7198 | -0.3317 | 0.0125 | 0.0000 | 0.0063 | UN |
| GA-2 | LA-2 | 4989909 | 0.0709 | 0.1647 | 0.1596 | 0.7780 | -0.5020 | 0.0118 | 0.0000 | 0.0059 | UN |
| GA-2 | OK | 4990661 | 0.0954 | 0.1230 | 0.1948 | 0.6765 | -0.2570 | 0.0120 | 0.0000 | 0.0060 | UN |
| GA-2 | MS-2 | 4990423 | 0.0914 | 0.1417 | 0.1920 | 0.7500 | -0.3381 | 0.0139 | 0.0039 | 0.0108 | UN |
| GA-2 | FL-3 | 4987887 | 0.1142 | 0.0881 | 0.2367 | 0.4026 | -0.1039 | 0.1410 | 0.0052 | 0.0757 | 4th |
| GA-2 | GA-1 | 4989918 | 0.1065 | 0.0902 | 0.2357 | 0.3999 | -0.1322 | 0.1453 | 0.0061 | 0.0788 | 4th |
| GA-2 | GA-3 | 4990155 | 0.1127 | 0.0757 | 0.2375 | 0.3486 | -0.0660 | 0.1484 | 0.0017 | 0.0759 | 4th |
| GA-2 | SC | 4988271 | 0.0855 | 0.1098 | 0.1894 | 0.4516 | -0.2495 | 0.0901 | 0.0000 | 0.0451 | 4th |
| GA-2 | NC | 4989165 | 0.0872 | 0.1112 | 0.1915 | 0.4648 | -0.2492 | 0.0673 | 0.0000 | 0.0337 | UN |
| GA-2 | MS-3 | 4989888 | 0.0828 | 0.1495 | 0.1749 | 0.7715 | -0.3885 | 0.0139 | 0.0000 | 0.0070 | UN |
| GA-2 | TXAR | 4990460 | 0.0614 | 0.1789 | 0.1459 | 0.8014 | -0.6147 | 0.0066 | 0.0000 | 0.0033 | UN |
| GA-2 | TX | 4989819 | 0.0718 | 0.1690 | 0.1651 | 0.7667 | -0.5255 | 0.0125 | 0.0045 | 0.0108 | UN |
| AL | LA-1 | 4990829 | 0.0912 | 0.1084 | 0.2218 | 0.6204 | -0.2501 | 0.1013 | 0.0000 | 0.0507 | 4th |
| AL | AR | 4990734 | 0.0920 | 0.1078 | 0.2229 | 0.6288 | -0.2449 | 0.0734 | 0.0046 | 0.0413 | UN |
| AL | MS-1 | 4991407 | 0.0981 | 0.0947 | 0.2314 | 0.5713 | -0.1747 | 0.0769 | 0.0015 | 0.0399 | UN |
| AL | LA-2 | 4991002 | 0.0871 | 0.1175 | 0.2268 | 0.6459 | -0.3143 | 0.0383 | 0.0000 | 0.0191 | UN |
| AL | OK | 4991600 | 0.1000 | 0.0983 | 0.2265 | 0.6106 | -0.1785 | 0.0219 | 0.0016 | 0.0125 | UN |
| AL | MS-2 | 4991286 | 0.0994 | 0.0991 | 0.2344 | 0.5994 | -0.1886 | 0.0554 | 0.0053 | 0.0330 | UN |
| AL | FL-3 | 4987682 | 0.0997 | 0.1209 | 0.2203 | 0.5512 | -0.2572 | 0.0635 | 0.0042 | 0.0360 | UN |
| AL | GA-1 | 4989139 | 0.0791 | 0.1446 | 0.1817 | 0.6281 | -0.4083 | 0.0507 | 0.0031 | 0.0284 | UN |
| AL | GA-3 | 4989475 | 0.0924 | 0.1359 | 0.2051 | 0.6030 | -0.3305 | 0.0867 | 0.0000 | 0.0434 | UN |
| AL | SC | 4988128 | 0.0799 | 0.1624 | 0.1935 | 0.6542 | -0.4967 | 0.0539 | 0.0000 | 0.0269 | UN |
| AL | NC | 4989016 | 0.0776 | 0.1503 | 0.1843 | 0.6289 | -0.4472 | 0.0390 | 0.0000 | 0.0195 | UN |
| AL | MS-3 | 4990982 | 0.0940 | 0.1031 | 0.2246 | 0.6102 | -0.2190 | 0.0420 | 0.0014 | 0.0224 | UN |
| AL | TXAR | 4991544 | 0.0731 | 0.1301 | 0.2002 | 0.6764 | -0.4270 | 0.0356 | 0.0000 | 0.0178 | UN |
| AL | TX | 4990790 | 0.0835 | 0.1337 | 0.2205 | 0.6840 | -0.3978 | 0.0337 | 0.0045 | 0.0214 | UN |
| LA-1 | AR | 4992670 | 0.1121 | 0.0467 | 0.2921 | 0.3678 | 0.0379 | 0.1370 | 0.0078 | 0.0763 | 4th |
| LA-1 | MS-1 | 4992843 | 0.1121 | 0.0581 | 0.2793 | 0.4569 | -0.0081 | 0.0622 | 0.0069 | 0.0380 | UN |
| LA-1 | LA-2 | 4992706 | 0.1023 | 0.0590 | 0.2845 | 0.4215 | -0.0340 | 0.1154 | 0.0013 | 0.0590 | 4th |
| LA-1 | OK | 4992983 | 0.1060 | 0.0542 | 0.2484 | 0.4474 | -0.0046 | 0.0449 | 0.0000 | 0.0225 | UN |
| LA-1 | MS-2 | 4993301 | 0.1213 | 0.0379 | 0.3082 | 0.3206 | 0.0885 | 0.1412 | 0.0096 | 0.0802 | 4th |
| LA-1 | FL-3 | 4988167 | 0.0914 | 0.1425 | 0.2023 | 0.7115 | -0.3563 | 0.0799 | 0.0014 | 0.0414 | UN |
| LA-1 | GA-1 | 4989981 | 0.0704 | 0.1695 | 0.1618 | 0.7983 | -0.5314 | 0.0196 | 0.0000 | 0.0098 | UN |
| LA-1 | GA-3 | 4990195 | 0.0774 | 0.1572 | 0.1696 | 0.7686 | -0.4436 | 0.0532 | 0.0000 | 0.0266 | UN |
| LA-1 | SC | 4989034 | 0.0709 | 0.1838 | 0.1716 | 0.7963 | -0.6127 | 0.0214 | 0.0026 | 0.0133 | UN |
| LA-1 | NC | 4989823 | 0.0718 | 0.1758 | 0.1720 | 0.7927 | -0.5712 | 0.0147 | 0.0000 | 0.0074 | UN |
| LA-1 | MS-3 | 4992838 | 0.1142 | 0.0469 | 0.2932 | 0.3817 | 0.0403 | 0.0843 | 0.0017 | 0.0439 | UN |
| LA-1 | TXAR | 4993691 | 0.0975 | 0.0571 | 0.2940 | 0.3997 | -0.0389 | 0.1253 | 0.0015 | 0.0641 | 4th |
| LA-1 | TX | 4992649 | 0.0915 | 0.0659 | 0.2527 | 0.4369 | -0.0889 | 0.1145 | 0.0000 | 0.0572 | 4th |
| AR | MS-1 | 4992820 | 0.1166 | 0.0441 | 0.2921 | 0.3686 | 0.0552 | 0.0910 | 0.0049 | 0.0504 | 4th |
| AR | LA-2 | 4992691 | 0.1028 | 0.0537 | 0.2843 | 0.3872 | -0.0100 | 0.0784 | 0.0059 | 0.0451 | 4th |
| AR | OK | 4992962 | 0.1115 | 0.0386 | 0.2635 | 0.3390 | 0.0642 | 0.1025 | 0.0021 | 0.0533 | 4th |
| AR | MS-2 | 4992829 | 0.1241 | 0.0439 | 0.3156 | 0.3682 | 0.0702 | 0.0918 | 0.0140 | 0.0599 | 4th |
| AR | FL-3 | 4987918 | 0.0899 | 0.1466 | 0.1972 | 0.7284 | -0.3725 | 0.0327 | 0.0000 | 0.0163 | UN |
| AR | GA-1 | 4989664 | 0.0729 | 0.1740 | 0.1676 | 0.8223 | -0.5416 | 0.0122 | 0.0000 | 0.0061 | UN |
| AR | GA-3 | 4989832 | 0.0809 | 0.1601 | 0.1775 | 0.7844 | -0.4461 | 0.0109 | 0.0000 | 0.0054 | UN |
| AR | SC | 4988688 | 0.0735 | 0.1924 | 0.1780 | 0.8353 | -0.6396 | 0.0134 | 0.0000 | 0.0067 | UN |
| AR | NC | 4989527 | 0.0724 | 0.1804 | 0.1725 | 0.8162 | -0.5860 | 0.0054 | 0.0000 | 0.0027 | UN |
| AR | MS-3 | 4992767 | 0.1215 | 0.0455 | 0.3160 | 0.3786 | 0.0605 | 0.0892 | 0.0123 | 0.0569 | 4th |
| AR | TXAR | 4993301 | 0.0921 | 0.0589 | 0.2712 | 0.4091 | -0.0597 | 0.0985 | 0.0029 | 0.0521 | 4th |
| AR | TX | 4992863 | 0.1059 | 0.0531 | 0.3023 | 0.3762 | -0.0006 | 0.1487 | 0.0121 | 0.0865 | 4th |
| MS-1 | LA-2 | 4993169 | 0.1005 | 0.0602 | 0.2634 | 0.4395 | -0.0415 | 0.0449 | 0.0054 | 0.0278 | UN |
| MS-1 | OK | 4993600 | 0.1156 | 0.0475 | 0.2647 | 0.4182 | 0.0374 | 0.0477 | 0.0042 | 0.0281 | UN |
| MS-1 | MS-2 | 4993437 | 0.1241 | 0.0465 | 0.3024 | 0.3997 | 0.0581 | 0.0656 | 0.0105 | 0.0433 | UN |
| MS-1 | FL-3 | 4988640 | 0.0965 | 0.1342 | 0.2069 | 0.6975 | -0.3051 | 0.0238 | 0.0013 | 0.0132 | UN |
| MS-1 | GA-1 | 4990575 | 0.0807 | 0.1537 | 0.1816 | 0.7575 | -0.4317 | 0.0228 | 0.0000 | 0.0114 | UN |
| MS-1 | GA-3 | 4990807 | 0.0892 | 0.1420 | 0.1920 | 0.7261 | -0.3518 | 0.0307 | 0.0019 | 0.0173 | UN |
| MS-1 | SC | 4989499 | 0.0750 | 0.1685 | 0.1749 | 0.7615 | -0.5199 | 0.0072 | 0.0000 | 0.0036 | UN |
| MS-1 | NC | 4990426 | 0.0809 | 0.1577 | 0.1888 | 0.7427 | -0.4602 | 0.0082 | 0.0000 | 0.0041 | UN |
| MS-1 | MS-3 | 4993396 | 0.1246 | 0.0460 | 0.3124 | 0.3915 | 0.0624 | 0.0591 | 0.0029 | 0.0324 | UN |
| MS-1 | TXAR | 4993737 | 0.0968 | 0.0673 | 0.2746 | 0.4688 | -0.0842 | 0.0436 | 0.0048 | 0.0266 | UN |
| MS-1 | TX | 4992956 | 0.1014 | 0.0721 | 0.2727 | 0.4929 | -0.0903 | 0.0480 | 0.0029 | 0.0269 | UN |
| LA-2 | OK | 4993310 | 0.0977 | 0.0653 | 0.2423 | 0.4890 | -0.0658 | 0.0438 | 0.0000 | 0.0219 | UN |
| LA-2 | MS-2 | 4993235 | 0.1069 | 0.0525 | 0.2840 | 0.3936 | 0.0037 | 0.0970 | 0.0071 | 0.0556 | 4th |
| LA-2 | FL-3 | 4988303 | 0.0806 | 0.1577 | 0.1868 | 0.7475 | -0.4588 | 0.0112 | 0.0000 | 0.0056 | UN |
| LA-2 | GA-1 | 4990126 | 0.0635 | 0.1844 | 0.1548 | 0.8237 | -0.6442 | 0.0000 | 0.0000 | 0.0000 | UN |
| LA-2 | GA-3 | 4990358 | 0.0691 | 0.1766 | 0.1595 | 0.8079 | -0.5651 | 0.0054 | 0.0000 | 0.0027 | UN |
| LA-2 | SC | 4989144 | 0.0655 | 0.2004 | 0.1692 | 0.8287 | -0.7408 | 0.0045 | 0.0000 | 0.0022 | UN |
| LA-2 | NC | 4990280 | 0.0684 | 0.1887 | 0.1755 | 0.8092 | -0.6746 | 0.0045 | 0.0000 | 0.0022 | UN |
| LA-2 | MS-3 | 4993169 | 0.1037 | 0.0538 | 0.2816 | 0.3972 | -0.0080 | 0.0716 | 0.0020 | 0.0378 | UN |
| LA-2 | TXAR | 4993795 | 0.0907 | 0.0695 | 0.2956 | 0.4481 | -0.1215 | 0.0738 | 0.0084 | 0.0453 | 4th |
| LA-2 | TX | 4992815 | 0.0850 | 0.0842 | 0.2521 | 0.5091 | -0.1977 | 0.0419 | 0.0000 | 0.0210 | UN |
| OK | MS-2 | 4993637 | 0.1170 | 0.0458 | 0.2678 | 0.4115 | 0.0458 | 0.0515 | 0.0017 | 0.0274 | UN |
| OK | FL-3 | 4988770 | 0.0973 | 0.1200 | 0.2006 | 0.6554 | -0.2452 | 0.0264 | 0.0038 | 0.0170 | UN |
| OK | GA-1 | 4990969 | 0.0864 | 0.1369 | 0.1886 | 0.7058 | -0.3445 | 0.0250 | 0.0000 | 0.0125 | UN |
| OK | GA-3 | 4991221 | 0.0941 | 0.1272 | 0.1966 | 0.6836 | -0.2797 | 0.0170 | 0.0000 | 0.0085 | UN |
| OK | SC | 4989906 | 0.0838 | 0.1552 | 0.1906 | 0.7277 | -0.4332 | 0.0114 | 0.0000 | 0.0057 | UN |
| OK | NC | 4990925 | 0.0865 | 0.1413 | 0.1956 | 0.6968 | -0.3711 | 0.0022 | 0.0000 | 0.0011 | UN |
| OK | MS-3 | 4993439 | 0.1171 | 0.0479 | 0.2752 | 0.4250 | 0.0391 | 0.0653 | 0.0080 | 0.0406 | UN |
| OK | TXAR | 4993983 | 0.0834 | 0.0569 | 0.2168 | 0.4126 | -0.0651 | 0.0594 | 0.0000 | 0.0297 | UN |
| OK | TX | 4993150 | 0.0949 | 0.0591 | 0.2388 | 0.4312 | -0.0474 | 0.0731 | 0.0031 | 0.0396 | UN |
| MS-2 | FL-3 | 4988729 | 0.0981 | 0.1330 | 0.2102 | 0.7002 | -0.2975 | 0.0520 | 0.0014 | 0.0274 | UN |
| MS-2 | GA-1 | 4990564 | 0.0797 | 0.1580 | 0.1782 | 0.7819 | -0.4485 | 0.0380 | 0.0033 | 0.0223 | UN |
| MS-2 | GA-3 | 4990886 | 0.0864 | 0.1498 | 0.1842 | 0.7698 | -0.3841 | 0.0259 | 0.0000 | 0.0130 | UN |
| MS-2 | SC | 4989469 | 0.0755 | 0.1772 | 0.1757 | 0.8000 | -0.5520 | 0.0154 | 0.0000 | 0.0077 | UN |
| MS-2 | NC | 4990417 | 0.0789 | 0.1659 | 0.1826 | 0.7829 | -0.4951 | 0.0091 | 0.0014 | 0.0060 | UN |
| MS-2 | MS-3 | 4993447 | 0.1274 | 0.0391 | 0.3205 | 0.3444 | 0.0938 | 0.0936 | 0.0140 | 0.0608 | 4th |
| MS-2 | TXAR | 4994159 | 0.0993 | 0.0521 | 0.2829 | 0.3745 | -0.0107 | 0.0809 | 0.0035 | 0.0440 | UN |
| MS-2 | TX | 4993044 | 0.1074 | 0.0638 | 0.2925 | 0.4465 | -0.0424 | 0.0807 | 0.0147 | 0.0550 | 4th |
| FL-3 | GA-1 | 4987829 | 0.0928 | 0.1133 | 0.2005 | 0.4764 | -0.2409 | 0.1180 | 0.0000 | 0.0590 | 4th |
| FL-3 | GA-3 | 4987779 | 0.1102 | 0.1048 | 0.2326 | 0.4575 | -0.1702 | 0.1370 | 0.0029 | 0.0714 | 4th |
| FL-3 | SC | 4986618 | 0.0903 | 0.1346 | 0.2034 | 0.5282 | -0.3351 | 0.0720 | 0.0000 | 0.0360 | UN |
| FL-3 | NC | 4987470 | 0.0948 | 0.1263 | 0.2131 | 0.5105 | -0.2923 | 0.0532 | 0.0000 | 0.0266 | UN |
| FL-3 | MS-3 | 4988197 | 0.0936 | 0.1433 | 0.2035 | 0.7361 | -0.3486 | 0.0235 | 0.0000 | 0.0118 | UN |
| FL-3 | TXAR | 4988757 | 0.0723 | 0.1748 | 0.1778 | 0.7812 | -0.5787 | 0.0206 | 0.0000 | 0.0103 | UN |
| FL-3 | TX | 4988206 | 0.0778 | 0.1596 | 0.1827 | 0.7253 | -0.4798 | 0.0491 | 0.0000 | 0.0246 | UN |
| GA-1 | GA-3 | 4990445 | 0.1015 | 0.0754 | 0.2282 | 0.3399 | -0.0904 | 0.1991 | 0.0049 | 0.1045 | 3rd |
| GA-1 | SC | 4990319 | 0.1037 | 0.0507 | 0.2641 | 0.2325 | 0.0047 | 0.2936 | 0.0071 | 0.1539 | 3rd |
| GA-1 | NC | 4990427 | 0.0980 | 0.0881 | 0.2428 | 0.3798 | -0.1558 | 0.1139 | 0.0027 | 0.0596 | 4th |
| GA-1 | MS-3 | 4990193 | 0.0717 | 0.1635 | 0.1615 | 0.7965 | -0.4954 | 0.0059 | 0.0000 | 0.0030 | UN |
| GA-1 | TXAR | 4990798 | 0.0571 | 0.1933 | 0.1486 | 0.8261 | -0.7470 | 0.0089 | 0.0000 | 0.0044 | UN |
| GA-1 | TX | 4990136 | 0.0663 | 0.1818 | 0.1660 | 0.7867 | -0.6388 | 0.0289 | 0.0000 | 0.0144 | UN |
| GA-3 | SC | 4989135 | 0.1002 | 0.0899 | 0.2358 | 0.3842 | -0.1516 | 0.1583 | 0.0047 | 0.0839 | 4th |
| GA-3 | NC | 4989868 | 0.0917 | 0.0987 | 0.2092 | 0.4158 | -0.1992 | 0.0799 | 0.0000 | 0.0399 | UN |
| GA-3 | MS-3 | 4990541 | 0.0857 | 0.1538 | 0.1869 | 0.7774 | -0.4079 | 0.0185 | 0.0015 | 0.0107 | UN |
| GA-3 | TXAR | 4990937 | 0.0624 | 0.1884 | 0.1532 | 0.8266 | -0.6695 | 0.0062 | 0.0000 | 0.0031 | UN |
| GA-3 | TX | 4990253 | 0.0714 | 0.1755 | 0.1688 | 0.7804 | -0.5659 | 0.0175 | 0.0000 | 0.0088 | UN |
| SC | NC | 4989610 | 0.0969 | 0.0840 | 0.2525 | 0.3492 | -0.1483 | 0.1427 | 0.0112 | 0.0825 | 4th |
| SC | MS-3 | 4989182 | 0.0749 | 0.1805 | 0.1785 | 0.8078 | -0.5790 | 0.0110 | 0.0014 | 0.0069 | UN |
| SC | TXAR | 4989765 | 0.0564 | 0.2107 | 0.1550 | 0.8365 | -0.8695 | 0.0061 | 0.0000 | 0.0031 | UN |
| SC | TX | 4989071 | 0.0635 | 0.2054 | 0.1669 | 0.8155 | -0.7819 | 0.0145 | 0.0000 | 0.0073 | UN |
| NC | MS-3 | 4990182 | 0.0764 | 0.1697 | 0.1804 | 0.7896 | -0.5265 | 0.0133 | 0.0000 | 0.0067 | UN |
| NC | TXAR | 4990657 | 0.0571 | 0.2005 | 0.1551 | 0.8241 | -0.8083 | 0.0040 | 0.0000 | 0.0020 | UN |
| NC | TX | 4989903 | 0.0648 | 0.1912 | 0.1683 | 0.7906 | -0.7064 | 0.0059 | 0.0000 | 0.0030 | UN |
| MS-3 | TXAR | 4993765 | 0.0965 | 0.0555 | 0.2817 | 0.3980 | -0.0327 | 0.0709 | 0.0000 | 0.0355 | UN |
| MS-3 | TX | 4992884 | 0.0995 | 0.0629 | 0.2731 | 0.4391 | -0.0566 | 0.0555 | 0.0051 | 0.0329 | UN |
| TXAR | TX | 4993436 | 0.0845 | 0.0773 | 0.2773 | 0.4744 | -0.1799 | 0.0689 | 0.0000 | 0.0345 | UN |

**Supplementary Table 4:** Potential recombination hotspots within the *A. mississippiensis* genome.

| Chromosome | Start | End | Width (Kb) | Rec. Rate (c/bp) | # Sites |
| --- | --- | --- | --- | --- | --- |
| 4 | 15109001 | 151699000 | 261 | 9.62e-9 | 5521 |
| 6 | 35280001 | 36652000 | 588 | 8.09e-9 | 1324 |

**Supplementary Table 5:** Enriched GO terms for genes found within putative recombination hotspots in *A. mississippiensis*.

|  | Fold Enrichment | FDR |
| --- | --- | --- |
| **GO biological process complete** |  |  |
| caveola assembly (GO:0070836) | 16.17 | 1.11E-02 |
| plasma membrane raft assembly (GO:0044854) | 14.37 | 1.73E-02 |
| membrane raft assembly (GO:0001765) | 12.93 | 2.35E-02 |
| plasma membrane raft organization (GO:0044857) | 11.76 | 3.20E-02 |
| semaphorin-plexin signaling pathway (GO:0071526) | 7.05 | 8.83E-04 |
| regulation of synaptic transmission, glutamatergic (GO:0051966) | 6.71 | 4.84E-02 |
| basement membrane organization (GO:0071711) | 6.68 | 2.45E-02 |
| response to organophosphorus (GO:0046683) | 4.99 | 1.56E-02 |
| regulation of actin filament depolymerization (GO:0030834) | 4.88 | 3.24E-02 |
| positive regulation of cellular process (GO:0048522) | 1.35 | 7.51E-03 |
| positive regulation of biological process (GO:0048518) | 1.31 | 1.50E-02 |
| biological regulation (GO:0065007) | 1.16 | 2.45E-02 |
| regulation of biological process (GO:0050789) | 1.15 | 4.42E-02 |
| **GO molecular function complete** |  |  |
| neuropilin binding (GO:0038191) | 0.64 | 5.66E-03 |
| semaphorin receptor binding (GO:0030215) | 10.89 | 1.08E-03 |
| chemorepellent activity (GO:0045499) | 8.62 | 9.99E-04 |
| protein binding (GO:0005515) | 6.09 | 3.88E-02 |
| **GO cellular component complete** |  |  |
| glutamatergic synapse (GO:0098978) | 2.17 | 2.11E-02 |
| cytoplasm (GO:0005737) | 1.18 | 4.00E-04 |
| membrane-bounded organelle (GO:0043227) | 1.14 | 1.95E-02 |
| organelle (GO:0043226) | 1.13 | 3.84E-03 |
| intracellular membrane-bounded organelle (GO:0043231) | 1.13 | 4.81E-02 |
| intracellular anatomical structure (GO:0005622) | 1.12 | 6.70E-04 |
| intracellular organelle (GO:0043229) | 1.12 | 2.48E-02 |
| cellular_component (GO:0005575) | 1.09 | 7.11E-10 |
| cellular anatomical structure (GO:0110165) | 1.09 | 5.52E-09 |

